# Nuclear wrinkles result from geometric adaptations to cellular and nuclear morphological changes in epithelial cells

**DOI:** 10.64898/2026.08.26.747252

**Authors:** Le Minh Khoa, Yohei Kono, Takeshi Shimi, Hiroshi Kimura

## Abstract

Mechanical cues influence cell behavior and fate and are frequently accompanied by changes in nuclear shape; however, how epithelial nuclei accommodate such deformations remains incompletely understood. Here, we investigated the formation and regulation of nuclear wrinkles (NWs), inward folds of the nuclear envelope, in human epithelial cells. Using quantitative confocal imaging in 2.5D spheroid cultures and controlled 2D monolayers, we found that NWs formed frequently in MCF10A cells but rarely in hTERT-RPE1 cells, indicating pronounced cell-type specificity. NW frequency increased with cell density and was tightly associated with coordinated geometric changes consistent with nuclear rounding. Disruption of F-actin organization, but not microtubules, robustly induced NW formation, and acute cell rounding triggered by trypsinization was sufficient to induce widespread wrinkling across multiple cell types. Live-cell imaging revealed that NWs are dynamic and reversible at low cell density but become stabilized under sustained confinement. NW formation occurred without detectable nuclear envelope rupture, DNA damage, or stress-associated histone phosphorylation. Quantitative analysis supports a passive geometric model in which redistribution of excess nuclear surface area accommodates nuclear shape remodeling, allowing epithelial nuclei to buffer mechanical constraints while preserving nuclear integrity.

## 1. Introduction

Mechanical stress from the extracellular and cellular microenvironments can regulate cell fate, behavior, and tissue homeostasis. As the largest and stiffest organelle, the nucleus that houses the genome acts as a major mechanical element within the cell (Kaminski et al., 2014; Maurer & Lammerding, 2019). Nuclear shape and mechanical features are affected by the integrated mechanical contributions of the cytoskeleton and nuclear structural components, whereby forces generated within the cytoskeleton are transmitted to the nucleus (Dahl et al., 2008; Isermann & Lammerding, 2013; Niethammer, 2021).

Among cytoskeletal systems, F-actin is the primary source of intracellular tension during interphase. Actomyosin contractility driven by non-muscle myosin II interactions with F-actin generates traction forces that regulate cell shape, adhesion, and migration (Chambliss et al., 2013; Khatau et al., 2009). Microtubules form a stiff cytoskeletal network that can bear compressive loads in living cells and thereby contribute to intracellular force balance (Brangwynne et al., 2006; Ju et al., 2024), while intermediate filaments, such as vimentin, limit nuclear deformation under large or sustained strain (Mendez et al., 2010; Patteson et al., 2019). Force transmission from the cytoskeleton to the nucleus is mediated primarily by linker of nucleoskeleton and cytoskeleton (LINC) complexes, which mechanically couple cytoskeletal filaments to the nuclear envelope through SUN–KASH interactions (Crisp et al., 2006; Lombardi et al., 2011). Beneath the inner nuclear membrane, the nuclear lamina forms a dense filamentous meshwork composed primarily of lamins, which are categorized into A- (lamin A and C) and B-types (lamin B1 and B2) (Shimi et al., 2008; Turgay et al., 2017). This lamina network provides structural support to the nuclear envelope, contributes to nuclear stiffness and shape stability, and serves as a key mechanical interface that distributes cytoskeleton-derived forces across the nuclear surface (Stephens et al., 2017).

In response to mechanical stress, the nucleus undergoes diverse deformations, including the formation of nuclear wrinkles (NWs), which have been reported in epithelial monolayers (Fricker et al., 1997; Johnson et al., 2003; Malhas et al., 2011) and three-dimensional spheroids (Jorgens et al., 2017), as well as in pathological contexts such as laminopathies (McClintock et al., 2006) and cancers (Bussolati et al., 2008; Wang, Dollahon, et al., 2024). However, the mechanical origin and biological significance of NW formation are not fully addressed.

NW formation is attributable to cytoskeleton-driven mechanisms, in which forces generated by cytoskeletal structures deform the nuclear envelope through physical coupling. In patterned endothelial cells and three-dimensional microenvironments, nuclear envelope indentations or wrinkles often coincide with actin and/or tubulin, suggesting an association between nuclear surface deformation and cytoskeletal reorganization (Cosgrove et al., 2021; Versaevel et al., 2014). The balance between microtubules and actin filaments appears to be important for maintaining nuclear morphology, as NWs increase when this balance is disturbed (Geng et al., 2023). However, despite these observations, direct evidence that individual cytoskeletal filaments locally indent the nuclear envelope to generate NWs remains limited. Cytoskeletal organization and cell adhesion may instead influence NW formation indirectly by altering cell and nuclear geometry.

Nuclear-intrinsic geometric models propose that NWs can arise under such conditions through passive folding of excess nuclear surface area, without requiring cytoskeletal indentation. Several studies have attributed NWs to nuclear volume reduction caused by cell rounding in fibroblasts (Kim et al., 2015) or by osmotic deswelling in developing fruit fly eggs (Jackson et al., 2023). A complementary framework emphasizes the nuclear drop model, in which nuclei across diverse human cancer cell lines possess intrinsic lamina surface area in excess of a sphere of equal volume, resulting in wrinkles in rounded nuclei and smoothing upon spreading, even when nuclear volume remains approximately constant (Dickinson et al., 2024; Wang, Abolghasemzade, et al., 2024). Thus, a unifying framework linking cytoskeletal and adhesion remodeling, cell and nuclear geometry, and NW formation remains lacking.

Here, we systematically investigate NW formation under controlled mechanical and geometric perturbations in epithelial cells. By combining quantitative imaging, targeted cytoskeletal perturbations, and live-cell analysis across multiple cell types, we show that NWs form reproducibly during both rapid and gradual changes in cell and nuclear geometry, including acute cell rounding, actin cytoskeleton disruption, and progressive confinement during long-term culture. NW formation under these conditions is not accompanied by global nuclear envelope rupture, detectable DNA damage, or stress-associated histone H3 Ser10 phosphorylation (H3S10ph), indicating that NWs do not inherently compromise nuclear or chromatin integrity. These findings suggest that NWs can arise as a mechanically accommodated nuclear shape change rather than as a manifestation of nuclear damage or acute stress.

## 2. Results

### 2.1 Nuclear wrinkles are cell-density dependent and cell-type specific

MCF10A cells form lumenized spheroids in 2.5D on-top Matrigel culture (Debnath et al., 2003; Fig. S1A). Compared to the early-stage spheroids (Day 4), many nuclei in late-stage spheroids (Day 10) showed intense nuclear wrinkles (NWs), typically depicted as signals of lamin B1 (LB1) within nuclear interior regions in z-series confocal projection images (Fig. S1B, C). Similar NW structures were also observed in confluent monolayer cells that spread outside the 2.5D matrix region in the same well, indicating that NWs are not exclusive to 3D environments and may be associated with geometric and mechanical constraints arising in high-density cellular environments (Fig. S1C, D). This observation led us to systematically compare how NWs emerge as cells transit from low to high density using a conventional 2D culture system.

MCF10A and hTERT-RPE1 (RPE1) were plated at the same cell density (1.5 × 10^4^ cells/cm^2^) on 2D plastic plates on Day 0, then fixed at Day 2, Day 4, and Day 6. The proportion of nuclei exhibiting NWs progressively increased as monolayers became denser in longer culture periods, although the extent was less obvious in RPE1 than in MCF10A cells (Fig. 1A). To quantify nuclear wrinkling, we detected intranuclear LB1 edge features by Canny-based edge detection and calculated a Wrinkling Index (WI; Cosgrove et al., 2021), defined as the fraction of LB1-positive edge pixels within the nuclear region (Fig. 1B; see Methods). MCF10A cells showed a strong density-dependent increase in WI, whereas RPE1 cells displayed minimal WI increase even under comparable high-density conditions (Fig. 1C). To characterize how nuclear geometry changes with density, we measured nuclear volume, nuclear cross-sectional area (the projected XY area of each nucleus), and nuclear height (thickness across z-axis confocal stack) (Fig. 1D–F). In MCF10A cells incubated for longer periods, nuclear volume decreased, accompanied by a decrease in nuclear cross-sectional area and a slight increase in nuclear height. Similar but milder changes were also observed in RPE1 cells. These coordinated geometric changes suggest that increasing cell density alters the mechanical environment of cells and promotes nuclear rounding, from a relatively flattened to a more spherical shape, which coincides with the emergence of NWs. Furthermore, analysis of a broader panel of cell lines revealed culture period- or cell density-dependent NW increases, similarly to MCF10A (Fig. S2A-E), reinforcing the view that NW formation is cell-type specific and linked to intrinsic nuclear geometry.

**Figure 1.**
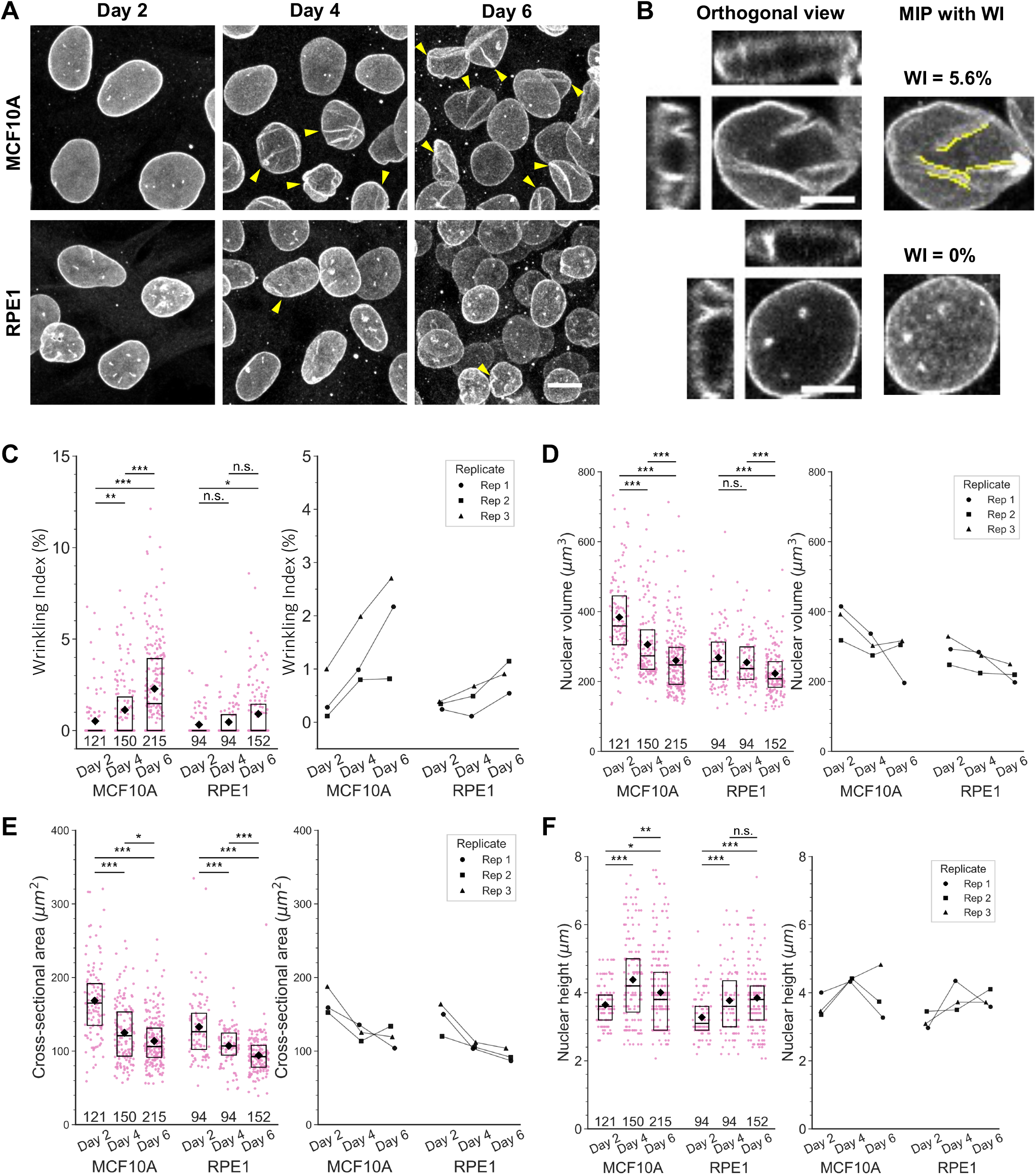
Nuclear wrinkle formation rate depends on cell density and cell type. **(A)** Representative images of LB1 in 2D monolayers of MCF10A (top) and hTERT-RPE1 (RPE1, bottom) cells at the indicated culture days. Maximum-intensity projection (MIP) images of confocal z-stacks are shown. Yellow arrowheads indicate nuclear wrinkles. **(B)** Example images of Wrinkling Index (WI) measurements for MCF10A (top) and RPE1 (bottom). Orthogonal views and MIP images with detected wrinkle edges (yellow) are shown. WI values are indicated. **(C)** Quantification of WI at different culture days. Left, WIs (%) of individual nuclei pooled from three biological replicates are shown as box plots. Numbers of nuclei are shown below the boxes. Box plots indicate the upper and lower quartiles with medians (lines) and means (diamonds). Individual data points are also plotted (magenta). Right, mean WIs for individual experiments are plotted, and values from the same set of experiments are connected. Statistical analysis was performed to assess day effects within each cell line using the Kruskal–Wallis test followed by Dunn’s multiple comparison test with Holm’s correction. *p < 0.05; **p < 0.01; ***p < 0.001; n.s., not significant. **(D–F)** Nuclear volume **(D)**, nuclear cross-sectional area **(E)**, and nuclear height **(F)** were quantified from the same dataset and analyzed as in **(C)**. Scale bars, 10 µm **(A)** and 5 µm **(B)**.

### 2.2 Inhibition of F-actin, but not microtubules, induces nuclear wrinkles

To investigate the relationship between cytoskeletal elements and NWs, we examined the localization of microtubules and F-actin relative to spontaneously occurring NWs by immunofluorescence in fixed cells. Some NWs partially coincided with microtubule and/or F-actin signals in both MCF10A and RPE1 cells, whereas others showed no apparent overlap (Fig. S3A–H). These observations suggest that cytoskeletal filaments may be associated with a subset of NWs, but do not establish that they directly generate them. We therefore next examined the effects of acute disruption of F-actin and microtubules on NW formation.

Cytoskeletal perturbations are also known to alter nuclear shape and mechanics (Makhija et al., 2016; Neelam et al., 2015). We therefore examined whether disrupting actin or microtubules influences NW formation using cells at relatively low density. MCF10A cells were treated with DMSO (control), 2 µM latrunculin B (LatB; an actin polymerization inhibitor), and 2 µM nocodazole (Noc; a microtubule polymerization inhibitor) for 30 minutes, before analyzing NW formation. Compared with the control, LatB induced prominent wrinkles, whereas Noc did not (Fig. 2A, B). Treatments with both LatB and Noc (LatBNoc) produced a phenotype similar to LatB alone, indicating that F-actin disruption is a primary driver of NW induction. Nuclear geometry analysis showed that LatB-treated and LatBNoc-treated nuclei exhibited decreased nuclear volume and cross-sectional area with increased nuclear height, indicating increased nuclear rounding (Fig. 2C–E).

**Figure 2.**
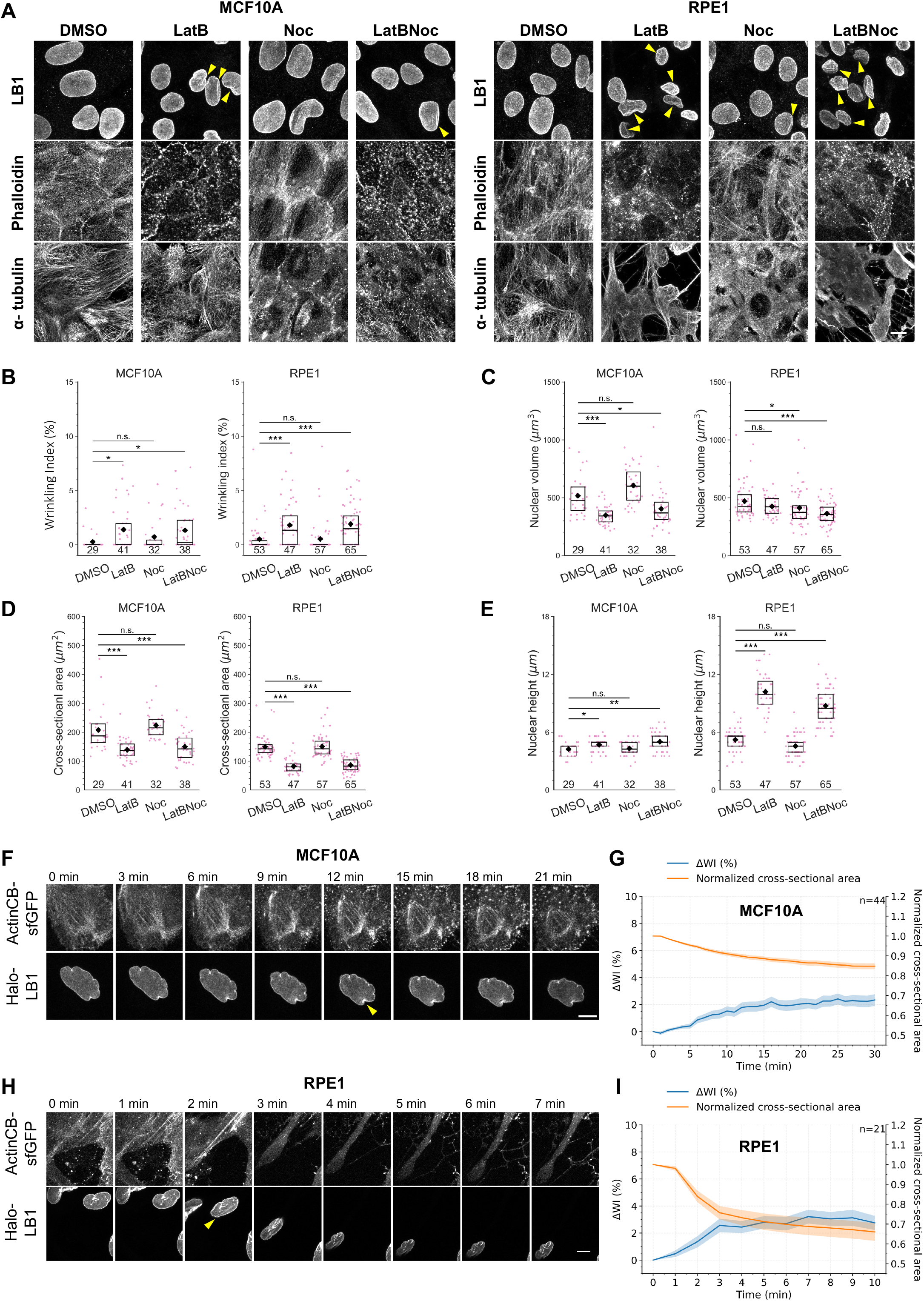
F-actin disruption, but not microtubule depolymerization, induces nuclear wrinkles. **(A)** Representative MIP images of confocal z-stacks of MCF10A (left) and RPE1 (right) cells treated for 30 min with DMSO (control), 2 µM latrunculin B (LatB), 2 µM nocodazole (Noc), or their combination (LatBNoc), and stained for LB1 (top), F-actin (phalloidin; middle), and α-tubulin (bottom). Yellow arrowheads indicate nuclear wrinkles. **(B–E)** Quantification of WI **(B)**, nuclear volume **(C)**, nuclear cross-sectional area **(D)**, and nuclear height **(E)** in cells treated with different cytoskeletal drugs. Data for MCF10A (left) and RPE1 (right), pooled from two biological replicates, are shown as box plots with statistical analysis (DMSO vs. each treatment), as in Fig. 1C–F. *p < 0.05; **p < 0.01; ***p < 0.001; n.s., not significant. **(F-I)** Time-lapse analysis in MCF10A (**F**, **G**) and RPE1 (**H**, **I**). **(F, H)** Representative time-lapse MIP images of cells stably expressing Halo-LB1 and ActinCB-sfGFP following administration of 2 µM LatB at t = 0. The same contrast scaling was applied to all frames. Arrows indicate the time point at which NWs first emerged. **(G, I)** Changes in WI relative to the first time point (ΔWI) and normalized nuclear cross-sectional area following LatB addition. Solid lines represent the mean, and shaded regions indicate ± SEM (standard error of mean). The number of cells analyzed is shown on each graph (MCF10A, n = 44; RPE1, n = 21). Scale bars, 10 µm.

RPE1 cells showed similar phenotypes to those of MCF10A, with increased NW formation following LatB and LatBNoc treatment (Fig. 2A, B). Whereas nuclear volume changed only subtly, nuclear height drastically increased (Fig. 2C–E), again consistent with nuclear rounding.

To further investigate the spatiotemporal dynamics of NWs during F-actin depolymerization, we performed time-lapse imaging of cells that stably express HaloTag-tagged LB1 (Halo-LB1) and superfolder GFP-tagged actin-specific chromobody (ActinCB-sfGFP, Rocchetti et al., 2014). When treated with LatB, nuclei shrank and NWs formed in both MCF10A and RPE1 cells, although the speed of their responses differed (Fig. 2F–I). Upon LatB administration, nuclear rounding and NW formation occurred gradually in MCF10A cells (Fig. 2F, G). In contrast, RPE1 cells responded rapidly, showing abrupt cell morphological changes (Fig. 2H, I). In both cell lines, the increase in WI was accompanied by a decrease in nuclear cross-sectional area (Fig. 2G, I), further supporting an association between nuclear rounding and NW formation.

Mouse embryonic carcinoma MC12 cells, human osteosarcoma U2OS cells, human cervical carcinoma HeLa cells, and human glioma NP-2 cells also formed NWs following LatB treatment (Fig. S4A–H), although the extent and dynamics of NW formation varied among cell lines. MC12 and U2OS cells exhibited relatively rapid and extensive NW formation, whereas NP-2, HeLa, and MCF10A cells were more gradual or less pronounced. These observations suggest that NW induction following F-actin disruption is conserved across multiple cell types, although both the kinetics and magnitude of the response vary among cell lines.

### 2.3 Cell rounding induces reversible nuclear wrinkle formation

LatB-induced F-actin depolymerization causes extensive cytoskeletal remodeling, including loss of cortical tension, altered intracellular force transmission, and changes in actomyosin contractility (Wakatsuki et al., 2001). To examine whether cell rounding alone, independently of direct cytoskeletal depolymerization, is sufficient to induce NWs, we next analyzed the effect of rapid cell rounding induced by trypsin treatment. Trypsinization has been reported to alter F-actin organization through disruption of cell-substrate adhesion and induction of cell rounding (Bereiter-Hahn et al., 1990; Sen & Kumar, 2009). Indeed, in cells treated with trypsin for 15 min, phalloidin staining showed a substantial loss of prominent actin stress fibers, although thin cortical protrusions and filopodia-like structures remained as cells became rounded (Fig. 3A). Thus, trypsinization induced extensive reorganization, rather than complete depolymerization, of the actin cytoskeleton. Although LatB and trypsin act through different primary mechanisms, both disrupted the spread cellular architecture, induced cell rounding, and were followed by nuclear rounding and NW formation.

**Figure 3.**
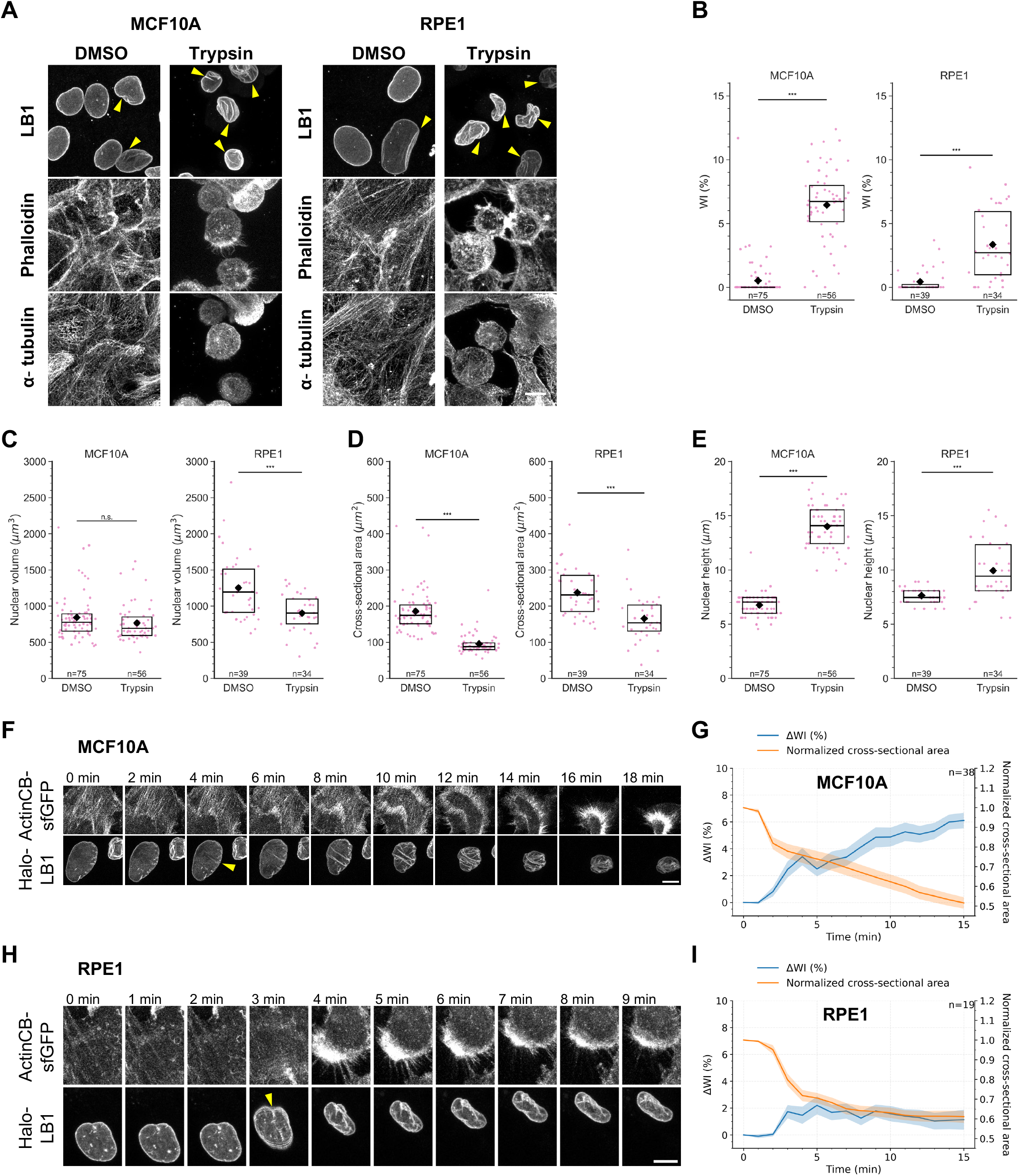
Trypsinization rapidly induces cell rounding and nuclear wrinkles. **(A)** Representative MIP images of confocal z-stacks of MCF10A (left) and RPE1 (right) cells treated for 15 min with DMSO (control) or trypsin (2.5 g/L) and stained for LB1 (top), F-actin (phalloidin; middle), and α-tubulin (bottom). Yellow arrowheads indicate nuclear wrinkles. **(B–E)** Quantification of WI **(B)**, nuclear volume **(C)**, nuclear cross-sectional area **(D)**, and nuclear height **(E)** in control and trypsinized cells. Data for MCF10A (left) and RPE1 (right) are shown as box plots and statistically analyzed using a two-sided Mann– Whitney U test. *p < 0.05; **p < 0.01; ***p < 0.001; n.s., not significant. Data were pooled from two biological replicates. **(F-I)** Time-lapse analysis in MCF10A (**F**, **G**) and RPE1 (**H**, **I**). **(F, H)** Representative time-lapse MIP images of MCF10A (**F**) and RPE1 (**H**) cells stably expressing Halo-LB1 and ActinCB-sfGFP following trypsin addition at t = 0 min. The same contrast scaling was applied to all frames. Arrows indicate the time point at which nuclear wrinkles (NWs) first emerged. **(G, I)** Time-course quantification of MCF10A (**G**) and RPE1 (**I**) nuclei, like those shown in **(F)** and RPE1 (**H**) cells, respectively. ΔWI and normalized nuclear cross-sectional area are shown as mean ± SEM. The number of cells analyzed is shown on each graph (MCF10A, n = 38; RPE1, n = 19). Scale bar, 10 µm.

Compared with the DMSO control, extensive NW formation was observed in both MCF10A and RPE1 cells following trypsinization, accompanied by nuclear rounding, as indicated by increased nuclear height and reduced nuclear cross-sectional area (Fig. 3B–E). Nuclear volume remained largely unchanged in MCF10A but was significantly reduced in RPE1 cells. Time-lapse imaging further demonstrated rapid and extensive NW formation following trypsinization in both MCF10A and RPE1 cells (Fig. 3F–I), as well as in U2OS, MC12, HeLa, and NP-2 cells (Fig. S5A–H), indicating that trypsinization-induced cell rounding is a general trigger for NW formation across multiple cell types.

To capture cell and nuclear morphological changes during trypsinization in detail, MCF10A and RPE1 cells that stably express Halo-LB1 and Lck-sfGFP, a plasma membrane-targeted fluorescent marker, were treated with a diluted trypsin solution to slow the reaction. Following trypsinization, both cell lines underwent rapid cell rounding, visualized using Lck-sfGFP, accompanied by nuclear deformation monitored by Halo-LB1 (Fig. 4A–D). Notably, changes in cell geometry consistently preceded the onset of NW formation. These observations support a model in which cell rounding drives subsequent nuclear rounding, which in turn leads to NW formation.

**Figure 4.**
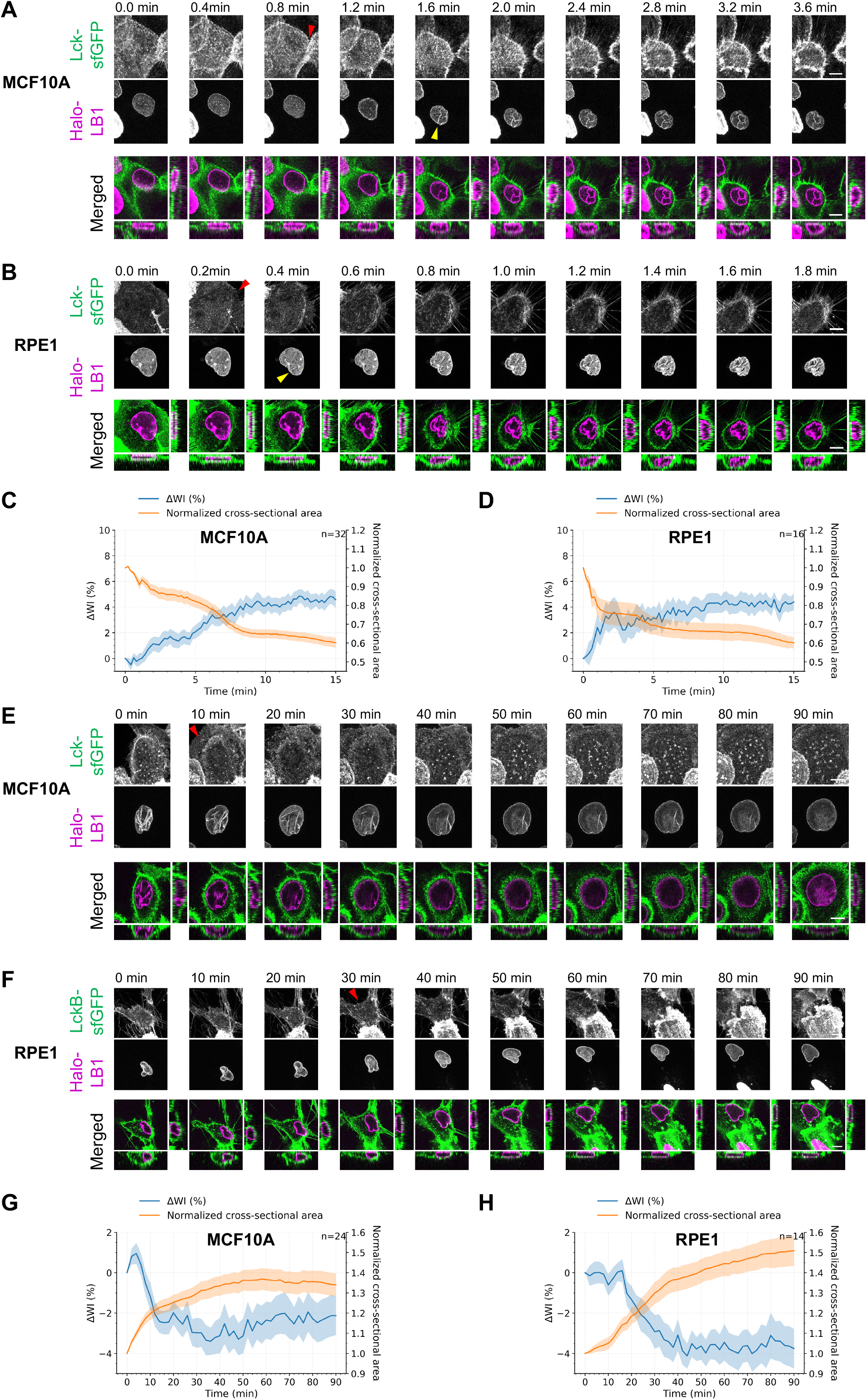
Cell rounding precedes nuclear wrinkle formation, and nuclear wrinkles are reversible upon cell spreading. **(A, B)** Representative time-lapse MIP (top and middle) and orthogonal-view (bottom) images of MCF10A **(A)** and RPE1 **(B)** cells stably expressing Halo-LB1 and Lck-sfGFP following treatment with diluted trypsin solution (1 g/L trypsin and 0.4 mmol/L EDTA) at t = 0. Red arrows indicate changes in plasma membrane geometry preceding the emergence of NWs, indicated by yellow arrowheads. **(C, D)** Time-course quantification of MCF10A **(C)** and RPE1 **(D)** nuclei following trypsinization. Changes in WI relative to the first time point (ΔWI) and normalized nuclear cross-sectional area are shown as mean ± SEM (MCF10A, n = 32; RPE1, n = 16). **(E, F)** Representative time-lapse MIP (top and middle) and orthogonal-view (bottom) images of trypsinized MCF10A **(E)** and RPE1 **(F)** cells stably expressing Halo-LB1 and Lck-sfGFP following replenishment with fresh culture medium to allow cell reattachment and spreading. Cell reattachments are observed (red arrowheads) earlier than NW deformation. **(G, H)** Time-course quantification of MCF10A **(G)** and RPE1 **(H)** nuclei during cell reattachment and spreading. ΔWI and normalized nuclear cross-sectional area are shown as mean ± SEM (MCF10A, n = 25; RPE1, n = 14). Scale bar, 10 µm.

To determine whether NW formation is reversible, the trypsin solution was gently replaced with fresh culture medium before complete cell detachment from the culture dish, allowing cells to reattach and spread. As the cells spread and restored a flattened morphology, their nuclei also flattened, as indicated by an increase in cross-sectional area, while NWs progressively diminished (Fig. 4E–H). These results indicate that NW formation is reversible and that restoration of a flattened cellular geometry alleviates nuclear wrinkling. As trypsinization may not be physiologically relevant, we also examined NWs in naturally rounded cell lines (KATOIII and Myeloma-653), which weakly attach to a dish surface. Remarkably, NWs were observed in nearly all nuclei (Fig. S6A, B), suggesting that rounded cells tend to have rounded nuclei with NWs.

### 2.4 Acute nuclear wrinkle formation is not accompanied by global nuclear envelope rupture

Because trypsinization and LatB treatment rapidly induced NW formation, we next examined whether this acute nuclear deformation is accompanied by a loss of nuclear envelope integrity. To this end, we expressed a superfolder Cherry-tagged nuclear localization signal (sfCherry-NLS) in MCF10A and RPE1 cells. Disruption of nuclear envelope integrity would permit sfCherry-NLS to leak into the cytoplasm. Despite robust NW formation following trypsinization, sfCherry-NLS remained predominantly localized to the nucleus in both cell lines (Fig. S7A–H). Quantitative analysis revealed only a modest decrease in the N/C ratio in a subset of MCF10A cells and little change in RPE1 cells. Similarly, no obvious leakage of sfCherry-NLS was detected following LatB treatment (Fig. S8A–H). In contrast, laser-induced nuclear envelope rupture caused a rapid and marked loss of nuclear NLS, resulting in a much greater decrease in the N/C ratio (Fig. S9A–D), consistent with the previous study (Kono et al., 2022). These results indicate that acute NW formation induced by trypsinization or LatB is not accompanied by widespread nuclear envelope rupture.

### 2.5 NW formation is dynamic and transient at low cell density without nuclear envelope rupture

Trypsinization represents an acute and artificial perturbation, distinct from the more gradual and sustained mechanical constraints associated with increasing cell density. We therefore examined NW dynamics and nuclear envelope integrity under more physiologically relevant conditions in untreated 2D cultures (Fig. 5A–D). Live-cell imaging of MCF10A cells that stably express Halo-LB1 revealed that, at low cell density (Day 2), NWs appeared transiently in association with fluctuations in nuclear shape during interactions with neighboring cells (Fig. 5A, C). In contrast, at high cell density (Day 5), NWs were more stable and long-lived (Fig. 5B, D). In either case, sfCherry-NLS remained confined to the nucleus without a detectable decrease in the N/C ratio during NW formation, indicating that nuclei maintain nuclear envelope integrity despite the formation of NWs (Fig. 5C, D). These observations suggest that nuclei can accommodate both transient geometric fluctuations and prolonged mechanical confinement without widespread nuclear envelope rupture.

**Figure 5.**
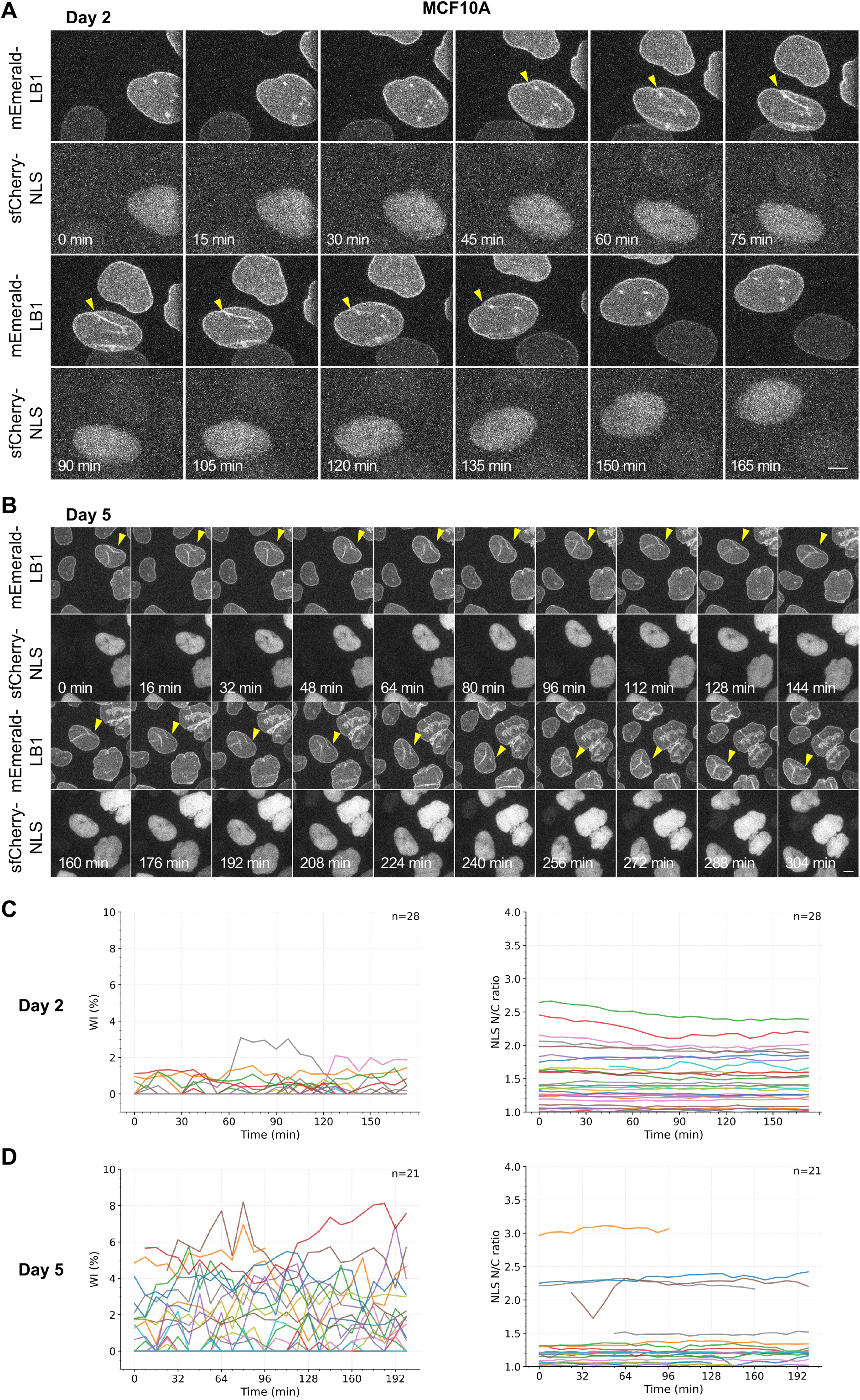
Nuclear wrinkles are dynamic and transient at low cell density without nuclear envelope rupture. **(A, B)** Representative time-lapse MIP images of MCF10A cells stably expressing Halo-LB1 and sfCherry-NLS at low cell density (Day 2) **(A)** and at high cell density (Day 5) **(B)**. Yellow arrowheads indicate nuclear wrinkles (NWs). **(C, D)** Time-course quantification of individual MCF10A cells at Day 2 **(C)** and Day 5 **(D)**. Left, WI of individual nuclei over time. Right, nuclear-to-cytoplasmic (N/C) sfCherry-NLS intensity ratio of individual cells over time. Each line represents a single cell (Day 2, n = 28; Day 5, n = 21). Scale bar, 10 µm.

To distinguish the effects of culture period from those of cell density, MCF10A cells that stably express Halo-LB1 and Lck-sfGFP were seeded at different densities on Day 0 and imaged on Day 4 (Fig. S10A–D). In low-density cultures, the establishment of cell–cell contact was accompanied by changes in cell morphology and nuclear shape, followed by transient NW formation (Fig. S10A, C). In contrast, cells surrounded by densely packed neighboring cells maintained altered cell and nuclear geometries, accompanied by longer-lasting NWs (Fig. S10B, D). These results suggest that local cell density, rather than culture duration per se, influences NW dynamics by altering cell morphology and nuclear shape.

### 2.6 Nuclear wrinkles are not associated with DNA damage or H3S10 phosphorylation

We next examined whether NWs are associated with histone H2AX Ser139 phosphorylation (γH2AX), a marker of DNA double-strand breaks (DSBs; Celeste et al., 2003; Hunt et al., 2013), or stress-induced histone H3 Ser10 phosphorylation (H3S10ph; Mahadevan et al., 1991; Soloaga et al., 2003). MCF10A and RPE1 cells cultured at different densities were fixed and stained with antibodies specific for γH2AX and H3S10ph. NW-positive nuclei showed no detectable increase in γH2AX (Fig. S11A, B) or H3S10ph (Fig. S11A, C).

As etoposide (ETP)-induced DNA damage has been reported to trigger nuclear deformation and invagination (Dellaire et al., 2009; Shokrollahi et al., 2024), we next investigated whether induction of DSBs promotes NW formation. MCF10A and RPE1 cells were treated with either vehicle control (DMSO) or 100 μM ETP for 1 h and then stained with γH2AX-specific antibody (Fig. S12A, B). Compared with the DMSO control, ETP treatment markedly increased γH2AX signals in both cell lines; however, no detectable increase in NW formation was observed. Consistent with these findings, time-lapse imaging of MCF10A and RPE1 cells for 1 h during ETP treatment showed no increase in NWs (Fig. S12C, D).

We further examined DNA damage under conditions that acutely induce NW formation by LatB or trypsin treatment. No detectable increase in γH2AX signals was observed in MCF10A or RPE1 cells treated with either LatB or trypsin (Fig. S13A–C). Together, these findings indicate that NW formation is neither associated with DNA damage nor stress-associated H3S10 phosphorylation, and that DNA double-strand breaks alone are insufficient to induce NW formation under the conditions tested in this study.

### 2.7 Nuclear wrinkles reflect a passive geometric adaptation of the nuclear envelope

Recent theoretical and experimental studies have proposed that nuclear shape is constrained by excess surface area (ESA) of the nuclear envelope and lamina (Alisafaei et al., 2019; Dickinson et al., 2024; Makhija et al., 2016). Consistent with this concept, all conditions that induced NW formation in the present study were accompanied by nuclear rounding, which reduces the surface area required to enclose a given nuclear volume. Following both LatB treatment and trypsinization, NW formation was accompanied by a decrease in nuclear cross-sectional area and an increase in nuclear height, indicating a transition from a flattened to a more rounded nuclear geometry (Figs. 2B–E and 3B–E). Notably, prominent NW formation occurred in trypsinized MCF10A cells despite little change in nuclear volume, indicating that nuclear volume reduction is not essential for NW formation. Moreover, the extent of NW formation differed markedly between MCF10A and RPE1 cells, suggesting that intrinsic differences in nuclear geometry may contribute to cell-type-specific susceptibility to NW formation.

To quantitatively assess this possibility, we estimated the relative ESA in taut nuclei with smooth nuclear surfaces (WI ≤ 0.1%) as the excess nuclear surface area normalized to the surface area of a sphere with the same nuclear volume (see Methods). Under low-density conditions, MCF10A nuclei exhibited significantly larger relative ESA than RPE1 nuclei (Fig. S14A), indicating that MCF10A nuclei possess a larger reservoir of excess nuclear surface area before exposure to conditions that induce nuclear rounding.

To further examine the relationship between ESA and NW formation, we analyzed relative ESA together with other nuclear geometric parameters using pooled cells, both without and with NWs, from all culture days under the same experimental conditions as in Fig. 1. Relative ESA negatively correlated with WI in both cell lines, although the strength of the correlation differed, being moderate in MCF10A (Pearson’s r = −0.42) and weak in RPE1 (Pearson’s r = −0.10) (Fig. S14B). In addition, relative ESA negatively correlated with nuclear height (Fig. S14C) and positively correlated with nuclear cross-sectional area (Fig. S14D) in both cell lines. These data suggest that, as nuclei become more rounded, excess nuclear surface is progressively accommodated as nuclear wrinkles, leading to a decrease in ESA and an increased WI. Consistently, WI was positively correlated with nuclear height and negatively correlated with nuclear cross-sectional area (Fig. S14E, F), with these correlations being more pronounced in MCF10A cells than in RPE1 cells. Together, these findings indicate that NW formation is closely associated with dynamic changes in excess nuclear surface area and nuclear geometry during nuclear rounding, supporting a passive geometric adaptation model.

## 3. Discussion

### 3.1 Mechanism of nuclear wrinkle formation

In this study, we demonstrate that nuclear wrinkles arise under both rapid and gradual changes in cell and nuclear geometry, including acute cell rounding, cytoskeletal perturbation, and increasing geometric constraints as cells become densely packed during prolonged culture. These observations suggest that NW formation is a general and robust nuclear response to geometric change.

Several mechanisms have been proposed to explain nuclear invaginations or wrinkles. These include direct mechanical impingement of cytoskeletal filaments onto the nucleus (Cosgrove et al., 2021; Versaevel et al., 2014), disruption of force balance between cytoskeletal networks (Geng et al., 2023), reduction in nuclear volume (Jackson et al., 2023; Kim et al., 2015), and passive surface buckling driven by excess nuclear membrane area (Dickinson et al., 2024). Our data provide partial support for nuclear volume-related mechanisms, as nuclear volume decreased under some conditions, including prolonged culture and acute LatB or trypsin treatment. However, the magnitude of these changes varied between cell types and treatments. Notably, trypsinized MCF10A cells formed prominent NWs despite little change in nuclear volume, indicating that a reduction in nuclear volume is not essential for NW formation. Following both LatB treatment and trypsinization, NW formation was accompanied by a decrease in nuclear cross-sectional area and an increase in nuclear height, consistent with nuclear rounding. Similarly, under gradual density-dependent conditions, increased cell density appears to influence NW formation, at least in part, by altering cell geometry and the mechanical environment of the nucleus. Our results are therefore most consistent with a passive geometric model in which the nuclear envelope contains excess surface area that is redistributed during nuclear shape changes under mechanical constraint. In this framework, NWs emerge as surface folds when the nuclear surface area exceeds the amount required to enclose the nuclear volume for a given shape. Both rapid nuclear shape transitions and slower confinement-driven remodeling can lead to such conditions, albeit on different timescales. Thus, NWs likely arise as a passive geometric adaptation of the nuclear envelope, reflecting the redistribution of excess surface area in response to changes in nuclear shape and mechanical constraint rather than requiring active force-driven invagination.

Consistent with this model, MCF10A nuclei exhibited a larger intrinsic relative ESA than RPE1 nuclei, which may contribute to their more frequent NW formation. A larger reservoir of excess nuclear surface area may make MCF10A nuclei more susceptible to wrinkling during density-associated nuclear rounding. In contrast, RPE1 cells responded more rapidly to acute LatB and trypsin treatments, suggesting that the kinetics of NW formation are additionally influenced by cell-type-specific mechanical properties. Several cellular and nuclear factors could contribute to these differences. Variations in cortical actomyosin organization and cell adhesion may affect the rate and extent of cell rounding and thereby alter the mechanical environment of the nucleus. Differences in cytoskeleton–nucleus coupling through the LINC complex may also influence how rapidly changes in cell geometry are transmitted to the nuclear envelope. In addition, nuclear-intrinsic properties, including the abundance and organization of nuclear lamins and chromatin, may affect nuclear stiffness, deformability, and susceptibility to wrinkling. These possibilities were not directly examined in the present study and therefore remain to be tested. Whereas these mechanical properties may influence the rapid response to acute perturbations, changes in cell morphology and local confinement under increasing cell-density conditions may contribute to the persistence of the rounded and wrinkled nuclear state. Together, our findings suggest that intrinsic ESA may influence the capacity for NW formation, whereas cell type-specific mechanical properties may shape its kinetics, extent, and persistence.

Although acute disruption of F-actin organization and cell–substrate adhesion induced cell rounding and NW formation, the endogenous remodeling of the cytoskeleton and cell adhesion during progressive monolayer crowding and spheroid maturation was not directly examined. In these settings, changes in cortical actomyosin organization, cell–cell adhesion, and cell–matrix adhesion may alter cell geometry and the forces acting on the nucleus. Determining how these processes develop spatially and temporally during culture growth will require direct imaging and perturbation of cytoskeletal and adhesion components in both 2D monolayers and spheroids. Further experiments that independently manipulate nuclear volume and surface area will also be required to clarify their respective contributions to nuclear shape and NW formation and to determine how the balance between nuclear volume and surface area interacts with other mechanical properties of the nucleus.

### 3.2 Potential functional roles of nuclear wrinkles

Even though our results are consistent with the view that NWs arise primarily as a passive and reversible adaptation to changes in nuclear geometry, this does not preclude functional roles for NWs in specific biological contexts. Indeed, it has been suggested that nuclear invaginations can participate in specialized cellular processes, such as neural activation-associated nuclear remodeling, which potentially facilitates signal-dependent nuclear organization (Feurle et al., 2020; Frey et al., 2023). Additionally, recent studies have shown that nuclear invaginations can promote DNA damage repair by increasing the local accessibility of repair machinery to damaged chromatin regions (Shokrollahi et al., 2024). However, under the experimental conditions in this study, NW formation was not associated with substantial nuclear envelope rupture, DNA double-strand breaks, or stress response-induced H3S10 phosphorylation, suggesting that NWs do not inherently compromise nuclear integrity and functional activities. NWs may therefore have context-dependent roles. In our in vitro cell culture systems, NWs might represent an intrinsic mechanical flexibility of the nucleus that accommodates changes in cell shape and cytoskeletal organization, particularly remodeling of the actin network, while helping preserve nuclear integrity. In contrast, when NWs persist under prolonged mechanical constraints in tissues, they may secondarily influence nuclear organization, signaling, or genome maintenance. Determining whether NWs are merely permissive structural adaptations or actively contribute to nuclear function will require future studies combining mechanical perturbations with direct readouts of nuclear function.

## 4. Methods

### 4.1 Cell culture and drug treatment

MCF10A cells (CRL-10317; ATCC) were cultured in DMEM/F-12 (FUJIFILM Wako) supplemented with 5% horse serum (New Zealand origin; Thermo Fisher Scientific), 20 ng/mL recombinant human EGF (AF-100-15; PeproTech), 10 μg/mL insulin, 0.5 μg/mL hydrocortisone (FUJIFILM Wako), 100 ng/mL cholera toxin from Vibrio cholerae (Sigma-Aldrich), and 1% glutamine–penicillin–streptomycin solution (GPS; Sigma-Aldrich). hTERT-RPE1 cells were cultured in DMEM/F-12 (FUJIFILM Wako) supplemented with 10% fetal bovine serum (FBS; qualified; Thermo Fisher Scientific) and 1% GPS. HeLa, U2OS, HEK293T, NP-2 and MC12 cells were cultured in DMEM, high glucose (Nacalai Tesque) supplemented with 10% FBS and 1% GPS. KATOIII cells were cultured in RPMI 1640 (Nacalai Tesque) supplemented with 10% FBS and 1% GPS. Mouse P3X63Ag8.653 (Myeloma-653) cells were cultured in GIT (Wako) supplemented with 10% FBS and 1% GPS. All cell lines were cultured at 37°C in a humidified atmosphere containing 5% CO□.

For cytoskeletal perturbation, cells were treated at 37°C for 30 min with 2 μM latrunculin B (LatB; FUJIFILM Wako), 2 μM nocodazole (Sigma-Aldrich) or a combination of both. For trypsinization, culture medium was replaced with 2.5 g/L trypsin-1 mmol/L EDTA solution (Nacalai Tesque) or a diluted solution containing 1 g/L trypsin-0.4 mmol/L EDTA for RPE1 cells. For etoposide treatment, cells were treated with 100 μM etoposide (ETP; Sigma-Aldrich) in the culture medium for 1 h at 37°C.

MCF10A cells were cultured in 2.5D using an on-top Matrigel method adapted from the published protocol (Debnath et al., 2003). Briefly, cells were trypsinized, centrifuged, and resuspended in the same culture medium used for 2D MCF10A culture, except that the horse serum concentration was reduced from 5% to 2%. The cell suspension was mixed with Matrigel to a final concentration of 2% Matrigel and 5 × 10^4^ cells/mL. Before seeding, wells of 8-well chamber slides were coated with a thin layer of Matrigel and allowed to solidify. A total of 400 μL of the cell-Matrigel suspension (2 × 10^4^ cells per well) was plated onto the Matrigel layer and incubated at 37°C with 5% CO□. The medium containing 2% Matrigel was replaced every 4 days.

### 4.2 Plasmid construction

PB533-EF1α-mEmerald-LB1-IRES-Neo (mEmerald-LB1) and its derivative PB533-EF1α-Halo-LB1-IRES-Puro (Halo-LB1) were constructed from the PB-EF1α-MCS-IRES-Neo PiggyBac transposon vector (PB533A-2; System Biosciences) using In-Fusion HD Cloning Kit (Takara Bio) in accordance with previously described materials and methods (Kono et al., 2022). The EF1α promoter in the PB-EF1α-MCS-IRES-Neo piggyBac transposon system vector (PB533A-2; System Biosciences) of Halo-LB1 was replaced with CMV early enhancer/chicken β-actin/rabbit β-globin (CAG) promoter by Ligation-Convenience Kit (Nippon Gene) and denoted as CAG-Halo-LB1. The pCDH-CMV-MCS-EF1α-Blast used for stable expression of the nuclear rupture marker NLS^SV40^-sfCherry-NLS^Myc^ (denoted as sfCherry-NLS) was described previously (Kono et al., 2022). The actin-specific chromobody sequence (Actin-Chromobody; Rocchetti et al., 2014) was cloned into the PB533A-2 vector containing superfolder GFP (sfGFP) using In-Fusion cloning system (Takara Bio) and denoted as ActinCB-sfGFP. The oligonucleotides used for this In-Fusion cloning were: 5’TACTCTAGAGCTAGCGAATTCCCATGGCTCAGGTGCAGCTG-3’ and 5’- TTTATGCATGGCCGCGGATCCGAACCTCCTCCACCGCTACCTC-3’.

To generate a plasma membrane marker, the N-terminal membrane-targeting sequence of Lck, corresponding to the first 10 amino acids, was synthesized as two complementary oligonucleotides. This peptide contains signals for co-translational myristoylation at Gly2 and palmitoylation of adjacent cysteine residues, which together mediate stable anchoring to the inner leaflet of the plasma membrane (Zlatkine et al., 1997). The complementary oligonucleotides were annealed, 5′-phosphorylated, and ligated into the PB533A-2 vector upstream of sfGFP, generating Lck-sfGFP. The oligonucleotide sequences were 5′- CTAGCCACCATGGGCTGCGTGTGCAGCAGCAACCCCGAG-3′ and 5′- CCGGCTCGGGGTTGCTGCTGCACACGCAGCCCATGGTGG-3′.

### 4.3 Transfection and stable cell line generation

Cells were seeded on 6-well plates one day before transfection to reach approximately 70% confluency at the time of transfection. Plasmids (0.5–1.0 μg total DNA per well) were transfected using Lipofectamine 3000 (Thermo Fisher Scientific) according to the manufacturer’s instructions. Stable cell lines were generated by cotransfection of a PB533-based plasmid with Super PiggyBac transposase vector (PB200PA-1; System Biosciences) at a 1:10 ratio. After replating cells on 10-cm dishes, cells were selected with 250–500 μg/mL G418 for 7 days or 0.25-1 μg/mL puromycin for 2–3 days. Cells that express fluorescent proteins were isolated using a SH800 cell sorter (Sony). For Halo-LB1 expression, MCF10A cells were transfected with either Halo-LB1 or CAG-Halo-LB1 while the other cell lines were transfected with CAG-Halo-LB1.

### 4.4 Lentiviral transduction

For stable introduction of the nuclear localization reporter sfCherry-NLS, a lentivirus-mediated transduction method was employed following the described protocol (Kono et al., 2022). Lentiviral particles were produced in HEK293T cells by co-transfecting the transfer plasmid encoding sfCherry-NLS together with packaging plasmids, including psPAX2 (plasmid #12260; Addgene) and pVSV-G (PT3343-5; Clontech), using Lipofectamine 3000 (Thermo Fisher Scientific).

At 48–72 hours post-transfection, viral supernatant was collected, filtered through a 0.45 μm filter, and applied to target cells in the presence of polybrene (hexadimethrine bromide; Sigma-Aldrich) to enhance transduction efficiency. After incubation, the medium was replaced with fresh culture medium, and cells were allowed to recover for 24–48 hours. Stable integration of the transgene was achieved through lentiviral genome insertion into the host genome. Cells that express fluorescent proteins were isolated using a SH800 cell sorter (Sony).

### 4.5 Immunofluorescence

Cells were fixed in 4% paraformaldehyde with 0.1% Triton X-100 in 250 mM HEPES-NaOH (pH 7.4) buffer (Pombo et al., 1998) for 5 min at room temperature (RT; ∼25°C). Fix solution was prepared using 16% Paraformaldehyde Aqueous Solution EM Grade (Electron Microscopy Sciences; 15710-S), 1 M HEPES-NaOH (pH7.4) (Nacalai Tesque), 10% Triton X-100 (Nacalai Tesque) by diluting into water (for Fig. 3A, Fig. S3, Fig. S12, Fig. S13, Fig. S15) or PBS (other figures). The effects of using different fix solutions on nuclear geometry and WI were evaluated separately (Fig. S15), demonstrating that they had minimal effects on WI and nuclear geometry.

After fixation, cells were washed with PBS, permeabilized with 1% Triton X-100 in PBS for 15 min at RT, followed by incubation with Blocking-One P solution (Nacalai Tesque) for 15 min at RT. Cells were incubated with primary antibodies in Can-Get-Signal immunostain Solution B (Solution B; TOYOBO) overnight at RT, followed by three washes with PBS, then incubated with secondary antibodies in Solution B for 1–2 h at RT. DNA was stained with Hoechst 33342 (1–2 μg/mL; Thermo Fisher Scientific). F-actin was visualized using Alexa Fluor 555–conjugated phalloidin (AF-555; 1:750–1:1000; Thermo Fisher Scientific). When combining with immunofluorescence, Hoechst 33342 and/or phalloidin were added in the secondary antibody mixture solution.

Primary antibodies used in this study were rabbit polyclonal anti-lamin B1 (1:500– 1:1000; 12987-1-AP; Proteintech), mouse monoclonal anti-α-tubulin (1:500–1:1000; 14-4502-82; Thermo Fisher Scientific), mouse monoclonal anti-γH2AX (Trakarnphornsombat and Kimura, 2023), and mouse monoclonal anti-H3S10ph (CMA311; Hayashi-Takanaka et al., 2009). The secondary antibodies used were donkey anti-mouse Ig (Jackson ImmunoResearch; 715-005-150), donkey anti-rabbit Ig (Jackson ImmunoResearch; 711-005-152), goat anti-mouse Ig (Jackson ImmunoResearch; 115-005-071), and goat anti-rabbit Ig (Jackson ImmunoResearch; 111-005-144), conjugated with Alexa Fluor 488 sulfodichlorophenol Ester (AF488; Thermo Fisher Scientific) or Cy5 N-hydroxysuccinimidyl Ester (Cytiva) (Hayashi-Takanaka et al., 2009; Trakarnphornsombat and Kimura, 2023).

Fluorescence images were collected using a point-scan confocal microscope (Ti2 with A1 system; Nikon) operated by the built-in software NIS-elements AR version 5.21.00 with an Apo TIRF 60× Oil DIC N2 (NA 1.49) objective lens (Nikon). Images were acquired using 405-nm, 488-nm, 561-nm and 640-nm laser lines (LU-N4; Nikon), GaAsP detectors, a 405/488/561/640 dichroic mirror, and 450/50, 525/50, 595/50, 700/75 emission filters with the following parameters: scan size, 512 × 512 pixels; scan speed, 1 frame/sec; line sequential scanning; zoom 1×.

### 4.6 Live-cell imaging

For figures other than Fig. 4A, 0.5 or 1.4 × 10^4^ cells were seeded on a glass-bottom 96-well plate (at a density of ∼1.5 or ∼4.5 × 10^4^ cells/cm^2^; AGC TechnoGlass). The following day, the culture plate was positioned on a heated chamber (Tokai Hit) maintained at 37°C and 5% CO_2_ on a point-scan confocal microscope (Ti2 with A1 system; Nikon). Images were acquired as above except zoom factor was set 1.5×.

For Fig. 4A, 1.4 × 10^5^ cells were seeded on a glass-bottom 35-mm dish (at a density of ∼1.5 × 10^4^ cells/cm^2^; AGC TechnoGlass). The following day, the culture dish was positioned on a heated chamber (Tokai Hit) maintained at 37°C and 5% CO_2_ on a spinning-disk confocal system (CSU-W1; Yokogawa) with an inverted microscope (Ti-E; Nikon), EM-CCD (iXon 3; Andor), a laser illumination system (LDI-NIR; Chroma Technology Japan), and a Plan Apo VC 100× Oil DIC N2 (NA 1.4) objective lens. Images were acquired using the built-in software NIS-elements ver. v5.11.03, with 470- and 555-nm laser lines, a 405/470/555/640 dichroic mirror, and 520/60 and 690/50 emission filters, with the following parameters: Z step, 0.5 µm; scan size, 1,024 × 1,024 pixels.

### 4.7 Image analysis

For fixed monolayer samples, nuclei were segmented in three dimensions based on the DNA-staining channel using Cellpose-SAM (Cellpose v4; Pachitariu et al., 2025). Segmentation masks were visually inspected and manually corrected, when necessary, using Napari. Nuclei touching the image borders and objects below a minimum volume threshold were excluded. Individual nuclei were cropped using their bounding boxes with a fixed margin for downstream analysis.

To quantify nuclear wrinkles, we adopted the previously reported Wrinkling Index (WI; Cosgrove et al., 2021) with minor modifications. LB1 signals within each nuclear crop were maximum-intensity projected using z-sections excluding the top and bottom planes. To select NW areas while excluding the nuclear rim, the nuclear mask was isotropically shrunk (scale factor 0.9), and an additional rim region (4 pixels) was excluded from analysis. LB1 images were normalized by percentile saturation (0.35%), and edge features were detected using a Canny edge detector (Gaussian smoothing σ = 1.5). An adaptive threshold was computed from the mean and standard deviation of the normalized LB1 intensity within the shrunk mask using threshold T = (mean − 1.5 × SD)/0.75, where the low and high Canny thresholds were set to T and 1.5T, respectively. Detected edge components were filtered by Feret diameter (≥ 2.5 µm) to remove small isotropic features. WI was defined as the percentage of detected edge pixels within the shrunk nuclear mask.

Nuclear geometric parameters were quantified from the manually corrected 3D segmentation masks. To account for anisotropic voxel dimensions, nuclear masks were rescaled along the z-axis to match the XY pixel size before three-dimensional geometric measurements. Nuclear volume was calculated from the number of voxels within the rescaled mask multiplied by the isotropic voxel volume. Nuclear surface area was estimated from the rescaled 3D binary mask by surface reconstruction using the marching cubes algorithm. Nuclear height was defined as the z-extent of the rescaled nuclear mask, and nuclear cross-sectional area was calculated from its XY projection. Relative excess surface area (ESA) was calculated as ESA = (A_measured_ − A_sphere_) / A_sphere_, where A_measured_ is the surface area of the reconstructed nuclear mask and A_sphere_ is the surface area of a sphere with the same volume as the measured nucleus. A_sphere_ was calculated as 4πr², where r is the radius of the equal-volume sphere. Thus, relative ESA represents the excess surface area of the reconstructed nuclear geometry relative to an equal-volume sphere.

Fine membrane surfaces within nuclear infoldings could not be fully resolved by confocal microscopy and were therefore only partially represented in the reconstructed nuclear masks. Consequently, the measured surface area and relative ESA of highly wrinkled nuclei may underestimate the actual nuclear envelope surface area. To compare intrinsic relative ESA between MCF10A and RPE1 cells, we restricted this analysis to taut nuclei with minimal wrinkling (WI ≤ 0.1%), for which nuclear surface geometry could be reconstructed more reliably. When nuclei with higher WI values were included in correlation analyses, relative ESA was interpreted as the measurable surface area of the reconstructed nuclear mask rather than as a complete estimate of the actual nuclear envelope surface area.

Image analysis of 3D spheroid samples followed the same general workflow as described above, with modifications to accommodate three-dimensional nuclear geometry and anisotropic voxel sampling.

For live-cell time-lapse datasets, nuclei were segmented for all frames using LB1 channel-intensity thresholding and manually corrected in Napari. Nuclear centroids were linked across frames using LapTrack, and only tracks spanning the full analysis window were included in time-course analyses. WI was computed for each nucleus at each time point from the LB1 maximum-intensity projection using the same general edge-detection procedure as described for fixed-cell images. Nuclear cross-sectional area was calculated from the segmented nuclear mask. For time-course analyses, WI was expressed as the change relative to the baseline frame (ΔWI = WI(t) − WI□), whereas nuclear cross-sectional area was normalized to the baseline value (A_norm_ = A(t)/A□).

To examine the spatial relationship between nuclear wrinkles (NWs) and cytoskeletal structures, fluorescence-intensity line profiles were obtained from representative confocal z-stacks using Fiji/ImageJ. For each NW, a one-pixel-wide line was manually drawn across the wrinkle in the selected central optical section and stored in the ROI Manager. Fluorescence-intensity profiles were measured along the same line in the LB1, α-tubulin, and phalloidin channels from the central optical section and neighboring z-sections. The line coordinates, z-position, pixel calibration, and raw fluorescence intensities were exported using a custom ImageJ macro. Representative image panels and line-profile plots were generated in Python. Fluorescence intensities were independently normalized to a range of 0–1 within each channel.

For analysis of nuclear envelope integrity, the nuclear masks and tracks generated as described above were used. The sfCherry-NLS channel was maximum-intensity projected along the z-axis for each time point. Mean nuclear NLS intensity was measured within the complete nuclear mask. Mean cytoplasmic NLS intensity was measured within a 5-pixel-wide ring outside the nuclear boundary, excluding regions overlapping other segmented nuclei. The nuclear-to-cytoplasmic NLS intensity ratio was calculated by dividing the mean nuclear sfCherry-NLS intensity by the mean intensity within the cytoplasmic ring.

### 4.8 Live-cell imaging and NE rupture induction by laser microirradiation

For NE rupture induction by laser microirradiation in Fig. S9, MCF10A and RPE1 cells stably expressing sfCherry-NLS and mEmerald-LB1 were placed in a heated chamber (Tokai Hit) maintained at 37°C and 5% CO on a point-scan confocal microscope (Ti2 with A1 system; Nikon). Fluorescence images were collected using the built-in NIS-Elements AR software version 5.21.00 with an Apo TIRF 60× Oil DIC N2 (NA 1.49) objective lens (Nikon). Images were acquired using 488- and 561-nm laser lines (LU-N4; Nikon), GaAsP detectors, a 405/488/561/640 dichroic mirror, and 525/50 and 595/50 emission filters with the following parameters: scan size, 256 × 256 pixels; scan speed, 1 frame/s; line-sequential scanning; and 10× zoom. After the first image was acquired, a 2.2 µm × 1.0 µm rectangular region was laser-microirradiated using a second scanner with the 405-nm laser at 100% transmission for 30 s, after which images were collected every 2 s.

For analysis of laser-induced nuclear envelope rupture, nuclei were segmented frame by frame from the mEmerald-LB1 channel using Li intensity thresholding after Gaussian smoothing (σ = 1.0). Morphological opening (radius, 1 pixel), removal of small objects, convex-hull filling, and hole filling were applied to generate complete nuclear masks. The masks were visually inspected and manually corrected using Napari. The mean sfCherry-NLS intensity was measured within the entire nuclear mask. Cytoplasmic sfCherry-NLS intensity was measured in a 5-pixel-wide ring outside the nuclear boundary, excluding regions overlapping other segmented nuclei. The nuclear-to-cytoplasmic (N/C) intensity ratio was calculated by dividing the mean nuclear sfCherry-NLS intensity by the mean intensity in the cytoplasmic ring.

### 4.9 Statistical analysis

Statistical analyses were performed using Python. For comparisons among multiple groups, the Kruskal–Wallis test was used followed by Dunn’s multiple comparison test with Holm’s correction. For pairwise comparisons, a two-sided Mann–Whitney U (MWU) test was applied.

## Acknowledgements

We thank members of the Kimura laboratory for experimental assistance and valuable scientific discussions, with special thanks to Mika Saotome for support with image analysis and Akito Ohi for constructing the PB533-EF1α-Halo-LB1-IRES-Puro vector. We also acknowledge the Bioscience Center, Integrative Bioscience Facility at the Institute of Science Tokyo for DNA sequencing services. One of the authors, Le MK, acknowledges financial support from the Otsuka Toshimi Scholarship Foundation for his PhD studies. ChatGPT (version 5.2) was used to improve the clarity of the manuscript. All AI-assisted suggestions were carefully reviewed, edited, and approved by the authors, who take full responsibility for the content.

## Declarations

### Author Contributions

**Le Minh Khoa:** Conceptualization, Formal Analysis, Investigation, Methodology, Software, Writing – Original Draft Preparation. **Yohei Kono:** Methodology, Resources, Writing – Review & Editing. **Takeshi Shimi:** Conceptualization, Methodology, Resources, Writing – Review & Editing. **Hiroshi Kimura:** Conceptualization, Funding Acquisition, Supervision, Visualization, Writing – Original Draft Preparation.

### Final Approval

All authors have reviewed the manuscript and agree to its submission.

### Ethics Approval

Not applicable.

### Data Availability

The original data are available upon request. All custom Python scripts for image analysis, data visualization, and statistical analysis, as well as representative example images, will be made publicly available on GitHub (https://github.com/Kimura-Lab/Le-et-al.-2026-).

