## Supplementary Figures for "Nuclear wrinkles result from geometric adaptations to cellular and nuclear morphological changes in epithelial cells"

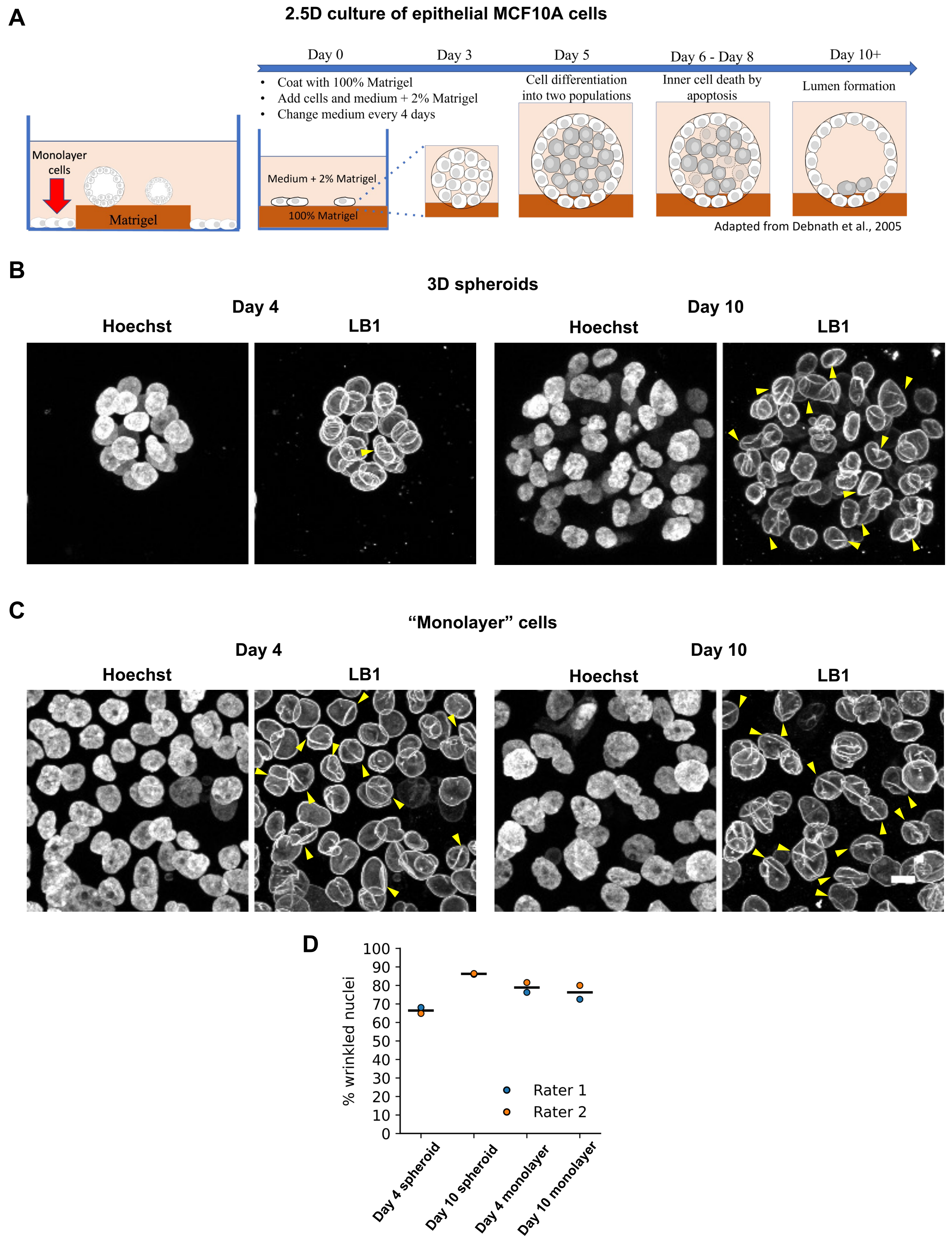


**Figure S1. 2.5D culture system and nuclear wrinkling in MCF10A cells.**
**(A)** Schematic illustration of the 2.5D culture setup for MCF10A cells. Epithelial spheroids and adjacent monolayer cells coexist within the same culture well.
**(B, C)** Representative confocal images of spheroids **(B)** and monolayer cells **(C)** from the same culture wells at Day 4 and Day 10, stained for DNA (Hoechst) and Lamin B1 (LB1). Yellow arrowheads indicate wrinkled nuclei.
**(D)** Percentage of wrinkled nuclei in spheroids and monolayers at Day 4 and Day 10. For each condition, the number of wrinkled nuclei was counted by two individuals (raters 1 and 2) and expressed as a percentage of the total number of nuclei. The horizontal line denotes the mean. Scale bars, 10 µm.

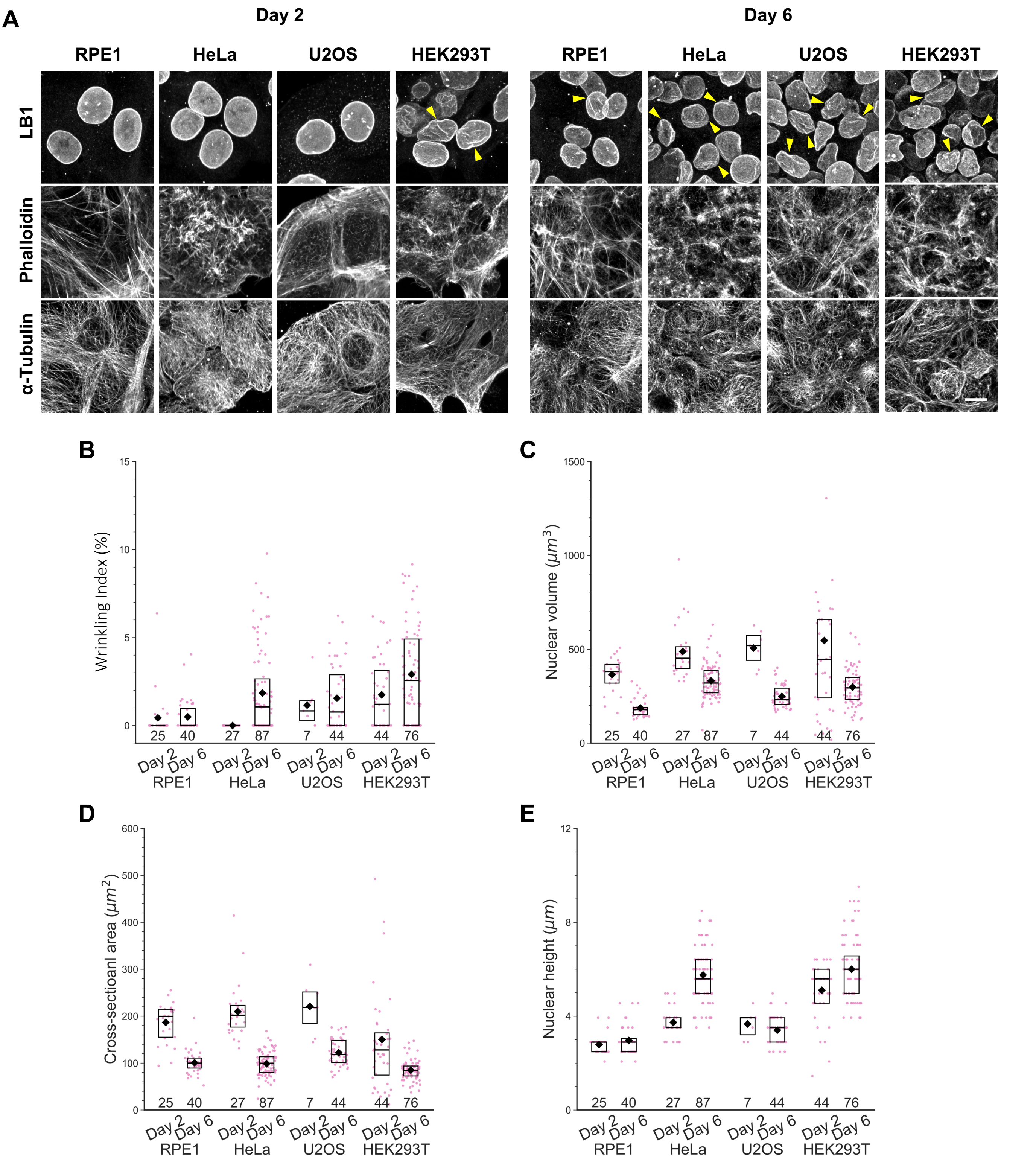


**Figure S2. Nuclear wrinkling and morphology in multiple human cell lines.**
**(A)** Representative immunofluorescence images of HEK293T, U2OS, HeLa, and RPE1 cells stained for Lamin B1 (LB1), F-actin (phalloidin), and α-tubulin. Yellow arrowheads indicate wrinkled nuclei.
**(B–E)** Quantification of nuclear wrinkling and morphology using cells like those shown in **(A)**. **(B)** Wrinkling Index (WI, %), **(C)** nuclear volume, **(D)** nuclear cross-sectional area, and **(E)** nuclear height. Box plots indicate the upper and lower quartiles with medians (lines) and means (diamonds). Individual data points are also plotted (magenta). Sample sizes (n) are shown below the boxes. Day 2 and Day 6 are shown for each cell line, based on one biological replicate.
Scale bars, 10 µm.

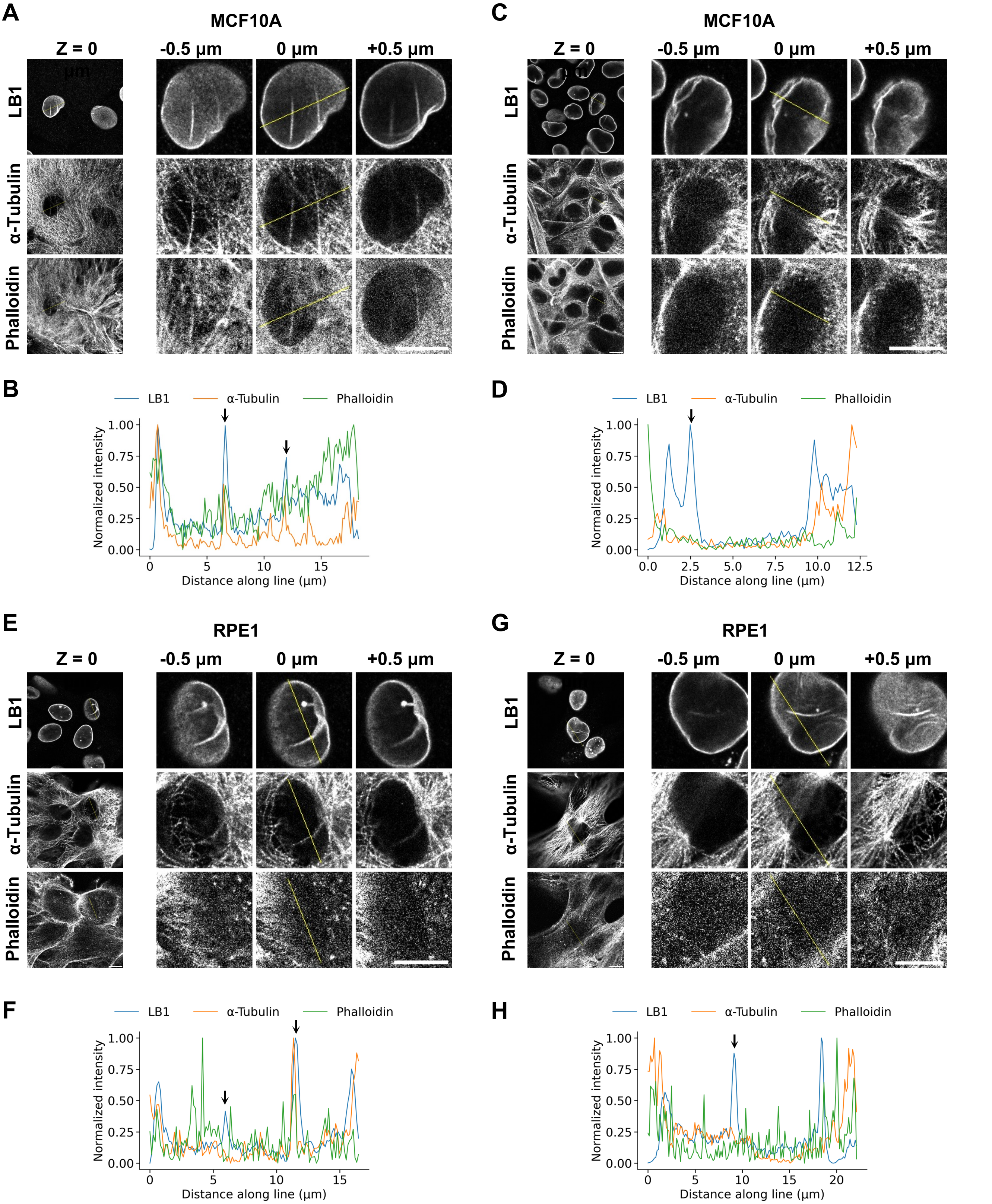


**Figure S3. Line-profile analysis of the spatial relationship between nuclear wrinkles and cytoskeletal structures.**
Representative confocal images and line profiles of MCF10A (**A–D**) and RPE1 (**E–H**) cells showing NWs.
(**A, C, E, G**) Confocal images. Left, central optical sections (0 µm). Right, enlarged views of the indicated nucleus at −0.5 µm, 0 µm, and +0.5 µm relative to the central optical section. The yellow line indicates the position used for line-profile analysis.
(**B, D, F, H**) Normalized fluorescence-intensity profiles of LB1, α-tubulin, and F-actin (phalloidin) measured along the line shown in **(A, C, E, G)**. Vertical arrows indicate the positions of the NWs showing LB1 peaks.
**(A, B, E, F)** Examples of cells containing NWs overlap with microtubules and/or F-actin.
**(C–H)** Examples of cells that containing NWs that do not overlap with microtubules and/or F-actin. In **E** and **F**, one NW does not overlap with either microtubules or F-actin, whereas another NW overlaps with microtubules.
Scale bars, 10 µm.

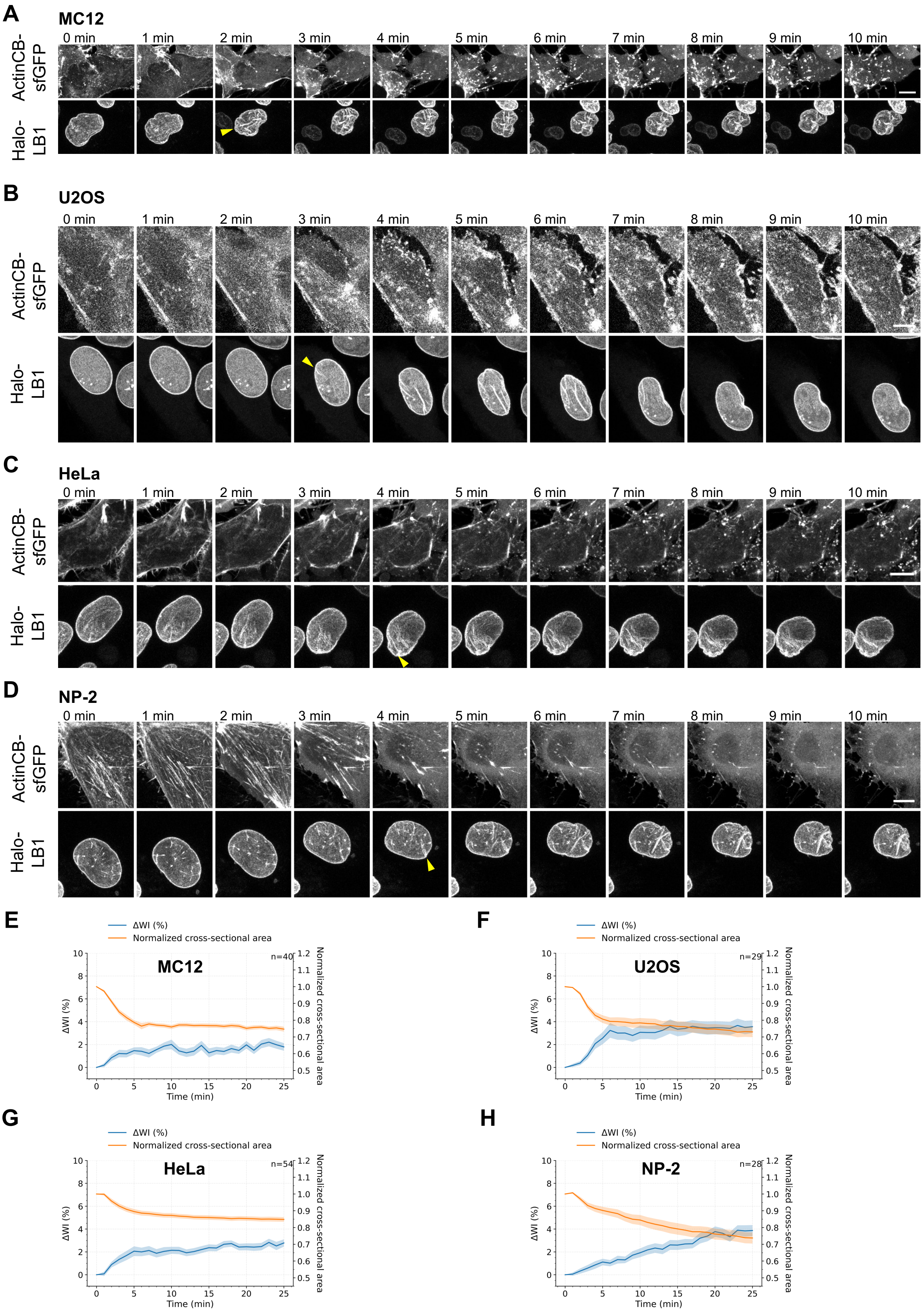


**Figure S4. Acute F-actin disruption induces nuclear wrinkling in U2OS and MC12 cells.**
**(A–D)** Representative time-lapse images of MC12 **(A)**, U2OS **(B)**, HeLa **(C)**, and NP-2 **(D)** cells following administration of 2 μM latrunculin B (LatB) at t = 0. Yellow arrowheads indicate the time point at which NWs first emerged.
**(E–H)** Changes in WI relative to the first time point (ΔWI) and normalized nuclear cross-sectional area of MC12 **(E)**, U2OS **(F)**, HeLa **(G)**, and NP-2 **(H)** cells following LatB addition. Means (solid line) ± standard errors of the mean (SEM; shaded regions) are shown, with sample sizes (n).
Scale bars, 10 μm.

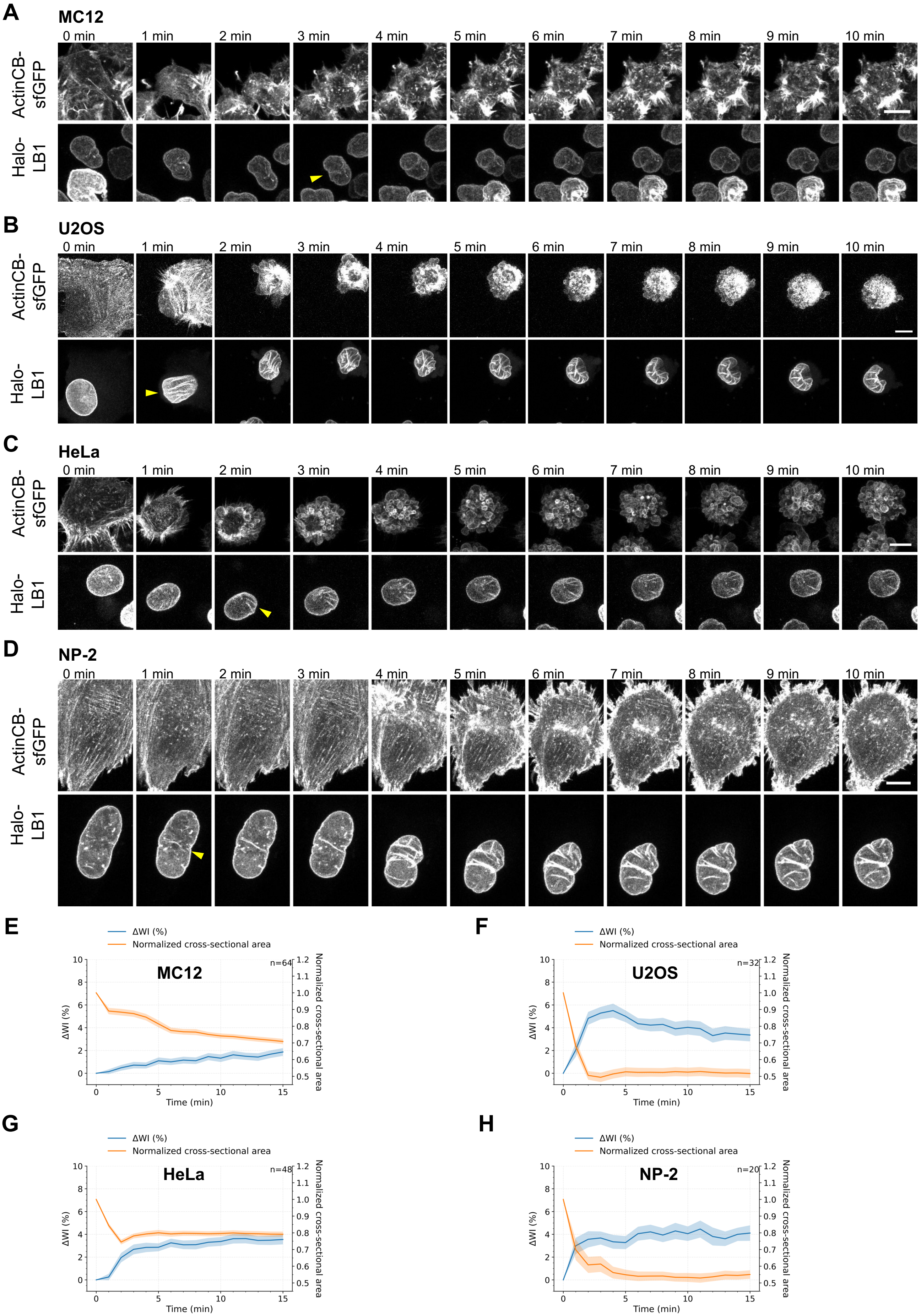


**Figure S5. Trypsinization induces dynamic nuclear wrinkling in multiple cell types.**
**(A–D)** Representative time-lapse images of MC12 **(A)**, U2OS **(B)**, HeLa **(C)**, and NP-2 **(D)** cells following trypsinization (2.5 g/L trypsin) at t = 0. Yellow arrowheads indicate the time point at which NWs first emerged.**(E–H)** Changes in WI relative to the first time point (ΔWI) and normalized nuclear cross-sectional area of MC12 **(E)**, U2OS **(F)**, HeLa **(G)**, and NP-2 **(H)** cells following trypsin treatment. Means (solid line) ± standard errors of the mean (SEM; shaded regions) are shown, with sample sizes (n).
Scale bars, 10 μm.

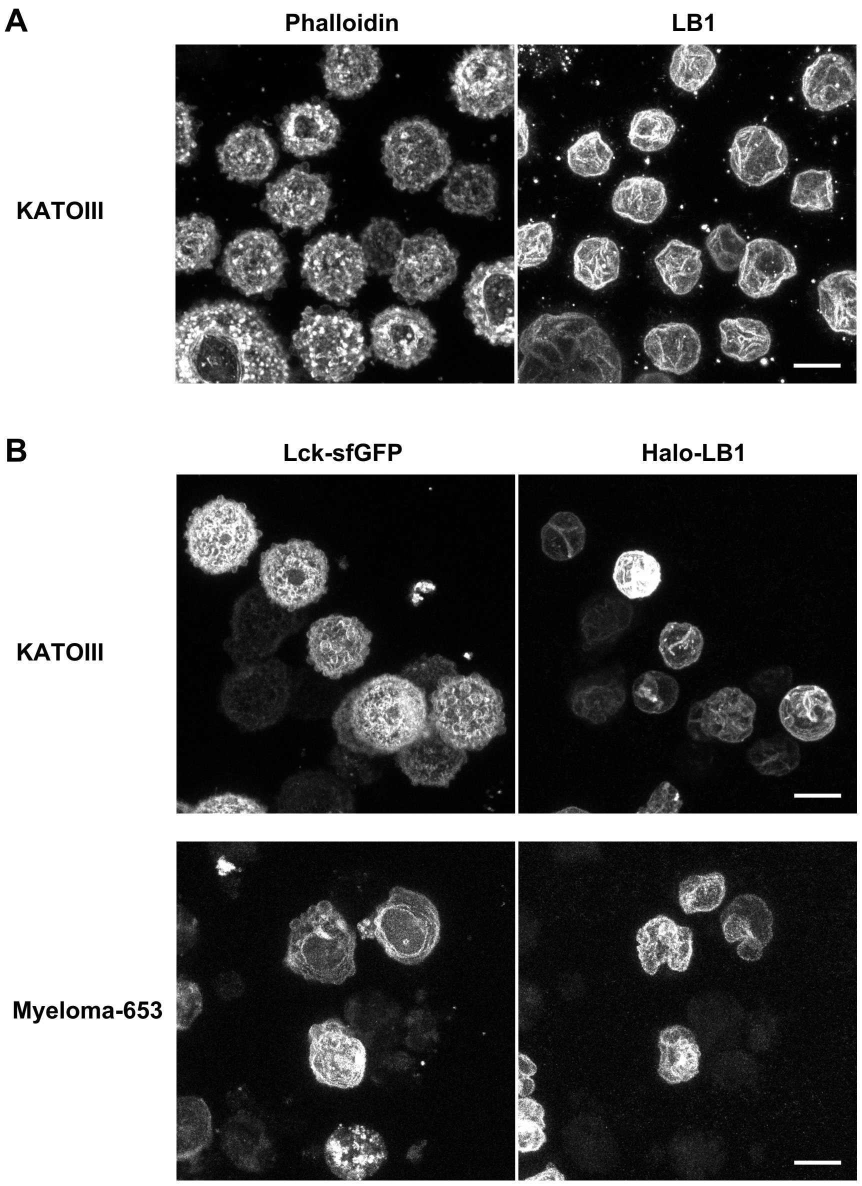


**Figure S6. Nuclear wrinkles in naturally non-adherent cell lines.**
(A) Representative MIP images of human KATOIII cells stained for phalloidin (left) and LB1 (right).
(B) Representative live-cell MIP images of KATOIII cells (top) and mouse P3X63Ag8.653 (Myeloma-653) cells (bottom) stably expressing Lck-sfGFP and Halo-LB1.
Scale bars, 10 µm.

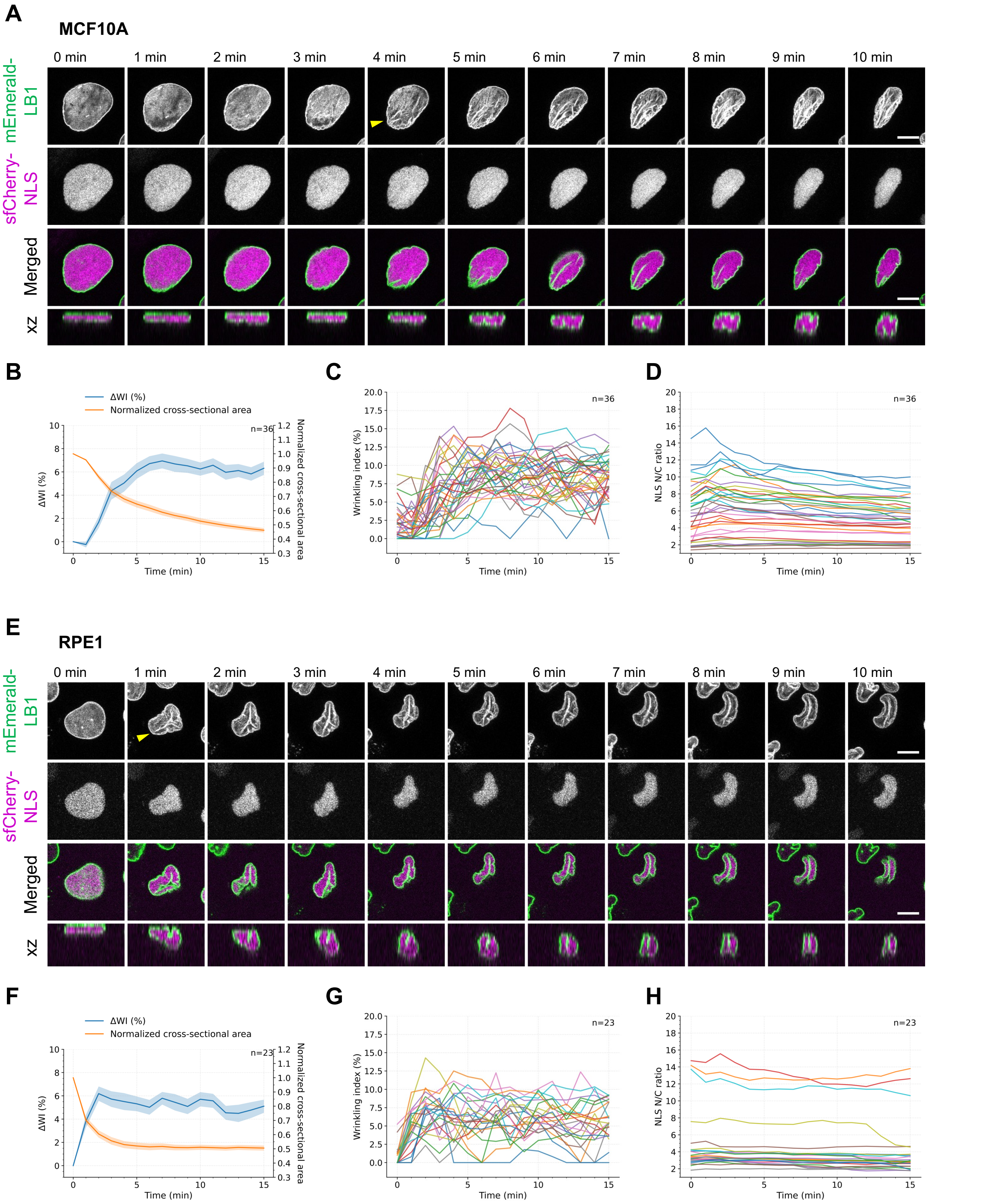


**Figure S7. Nuclear wrinkle formation during trypsinization is not accompanied by global nuclear envelope rupture.**

**(A)** Representative MIP and orthogonal-view images of MCF10A cells expressing mEmerald-LB1 and sfCherry-NLS following trypsinization (1 g/L trypsin) at t = 0.

**(B–D)** Quantitative analysis of MCF10A cells. **(B)** Changes in ΔWI and normalized nuclear cross-sectional area (mean ± SEM, n = 36). **(C, D)** Time courses of WI **(C)** and the nuclear-to-cytoplasmic intensity ratio of sfCherry-NLS (NLS N/C ratio) **(D)** in individual cells.

**(E)** Representative MIP and orthogonal-view images of RPE1 cells expressing mEmerald-LB1 and sfCherry-NLS following trypsinization (1 g/L trypsin) at t = 0.

**(F–H)** Quantitative analysis of RPE1 cells. **(F)** Changes in ΔWI and normalized nuclear cross-sectional area (mean ± SEM, n = 23). **(G, H)** Time courses of WI **(G)** and NLS N/C ratio **(H)** in individual cells.

Yellow arrowheads indicate the time point at which NWs first emerged. Scale bars, 10 µm.

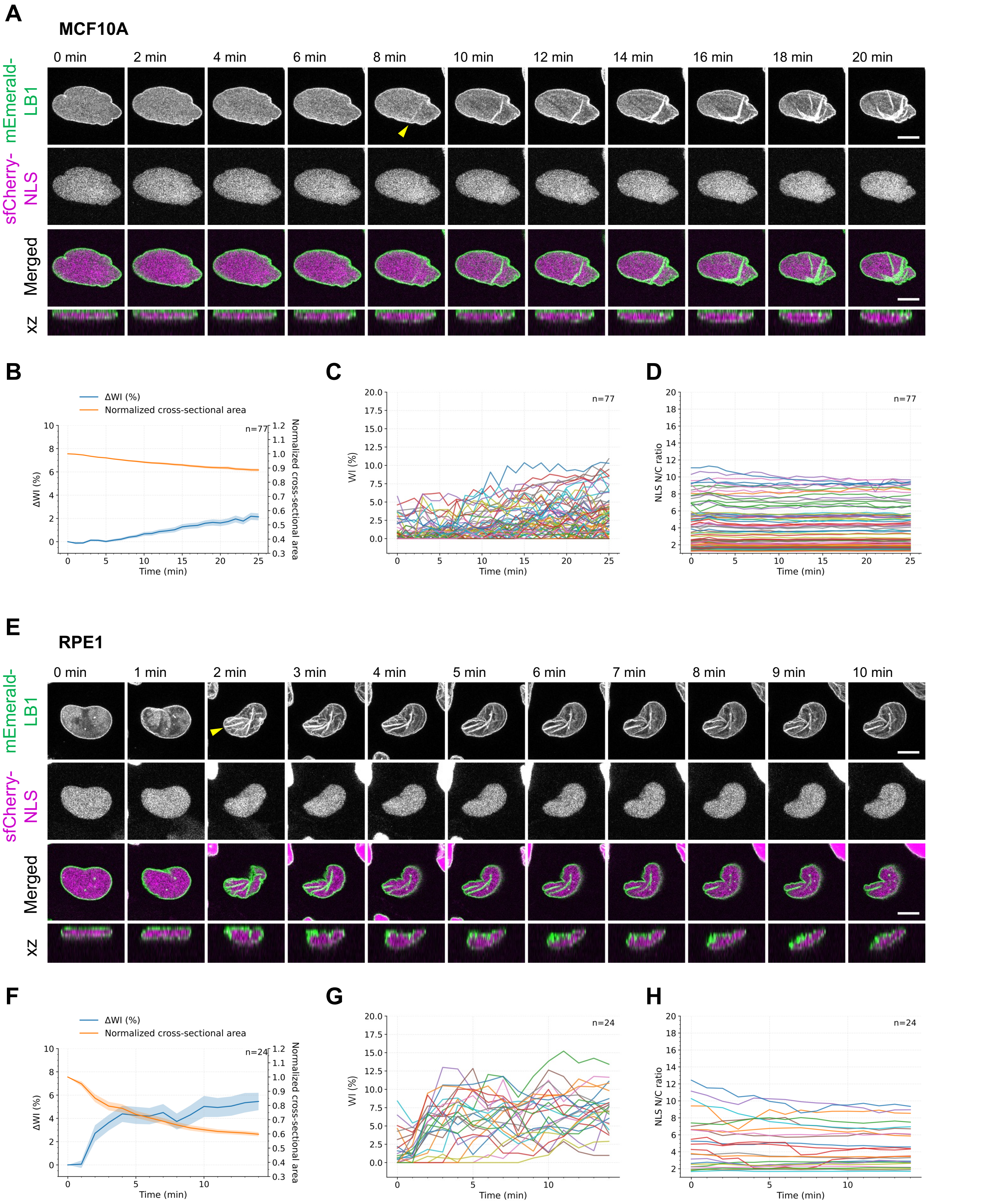


**Figure S8. Nuclear wrinkle formation following LatB treatment is not accompanied by global nuclear envelope rupture.**
**(A)** Representative MIP and orthogonal-view images of MCF10A cells expressing mEmerald-LB1 and sfCherry-NLS following treatment with 2 μM LatB at t = 0.
**(B–D)** Quantitative analysis of MCF10A cells. **(B)** Changes in ΔWI and normalized nuclear cross-sectional area (mean ± SEM, n = 77). **(C, D)** Time courses of WI **(C)** and NLS N/C ratio **(D)** in individual cells.
**(E)** Representative MIP and orthogonal-view images of RPE1 cells expressing mEmerald-LB1 and sfCherry-NLS following treatment with 2 μM LatB at t = 0.
**(F–H)** Quantitative analysis of RPE1 cells. **(F)** Changes in ΔWI and normalized nuclear cross-sectional area (mean ± SEM, n = 24). **(G, H)** Time courses of WI **(G)** and NLS N/C ratio **(H)** in individual cells.
Yellow arrowheads indicate the time point at which nuclear wrinkles (NWs) first emerged. Scale bars, 10 μm.

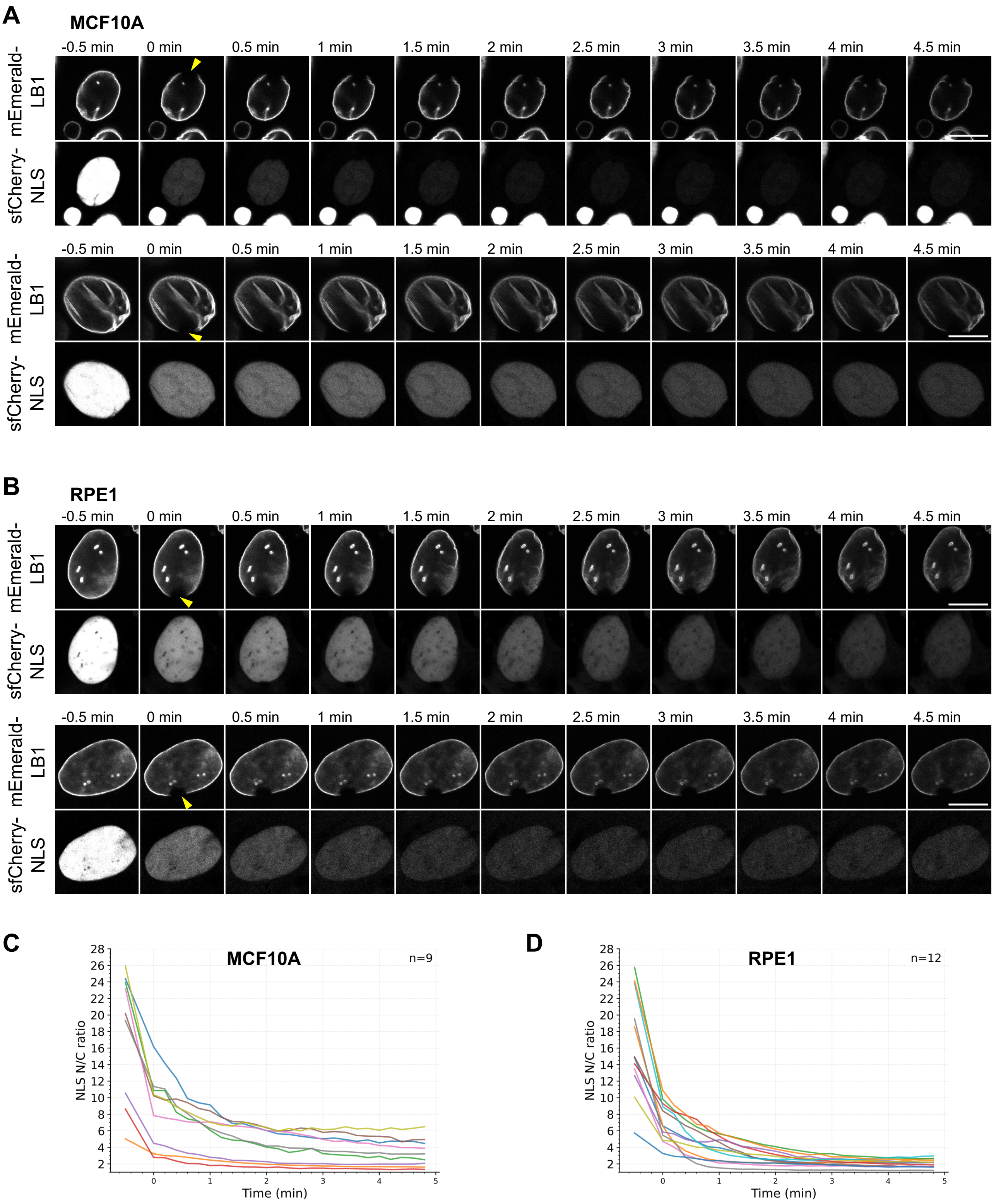


**Figure S9. Laser-induced nuclear envelope rupture causes rapid loss of nuclear NLS.**

**(A, B)** Representative confocal images of MCF10A **(A)** and RPE1 **(B)** cells stably expressing mEmerald-LB1 and sfCherry-NLS following laser-induced nuclear envelope rupture. Yellow arrowheads indicate the time point at which nuclear envelope rupture occurred.

**(C, D)** Time-resolved changes in NLS N/C ratio are shown for individual MCF10A **(C**, n = 9**)** and RPE1 **(D**, n = 12**)** cells.

Scale bars, 10 μm.

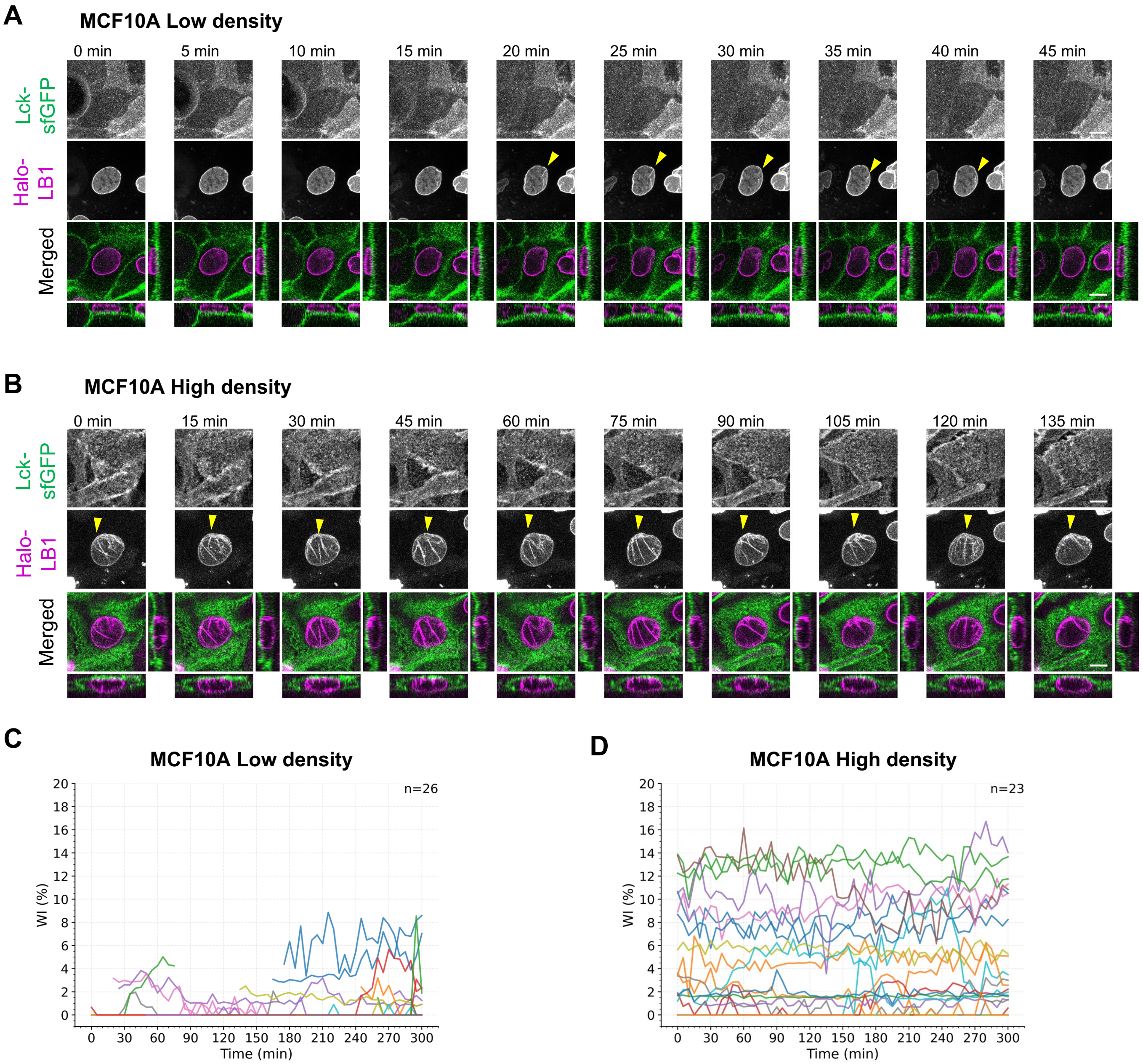


**Figure S10. Nuclear wrinkle formation is dynamic and transient at low cell density.**
**(A, B)** Time-lapse maximum-intensity projection (MIP) and orthogonal-view images of representative MCF10A cells stably expressing Lck-sfGFP and Halo-LB1 cultured at low density **(A**; $1.5 \times 10^4$ cells/cm² seeded at Day 0; imaged at Day 4; 5-min intervals**)** or high density (**B**; $4.5 \times 10^4$ cells/cm² seeded at Day 0; imaged at Day 4; 15-min intervals**)**. **(A)** At low density, NW formation was transient and occurred following cell-cell contact from the left side. **(B)** At high density, NWs were persistently observed. Yellow arrowheads indicate the appearance of NWs.
**(C, D)** Time-resolved changes in WI of individual nuclei in low-density (**C**, n = 26) and high-density (**D**, n = 23) cultures.
Scale bars, 10 µm.

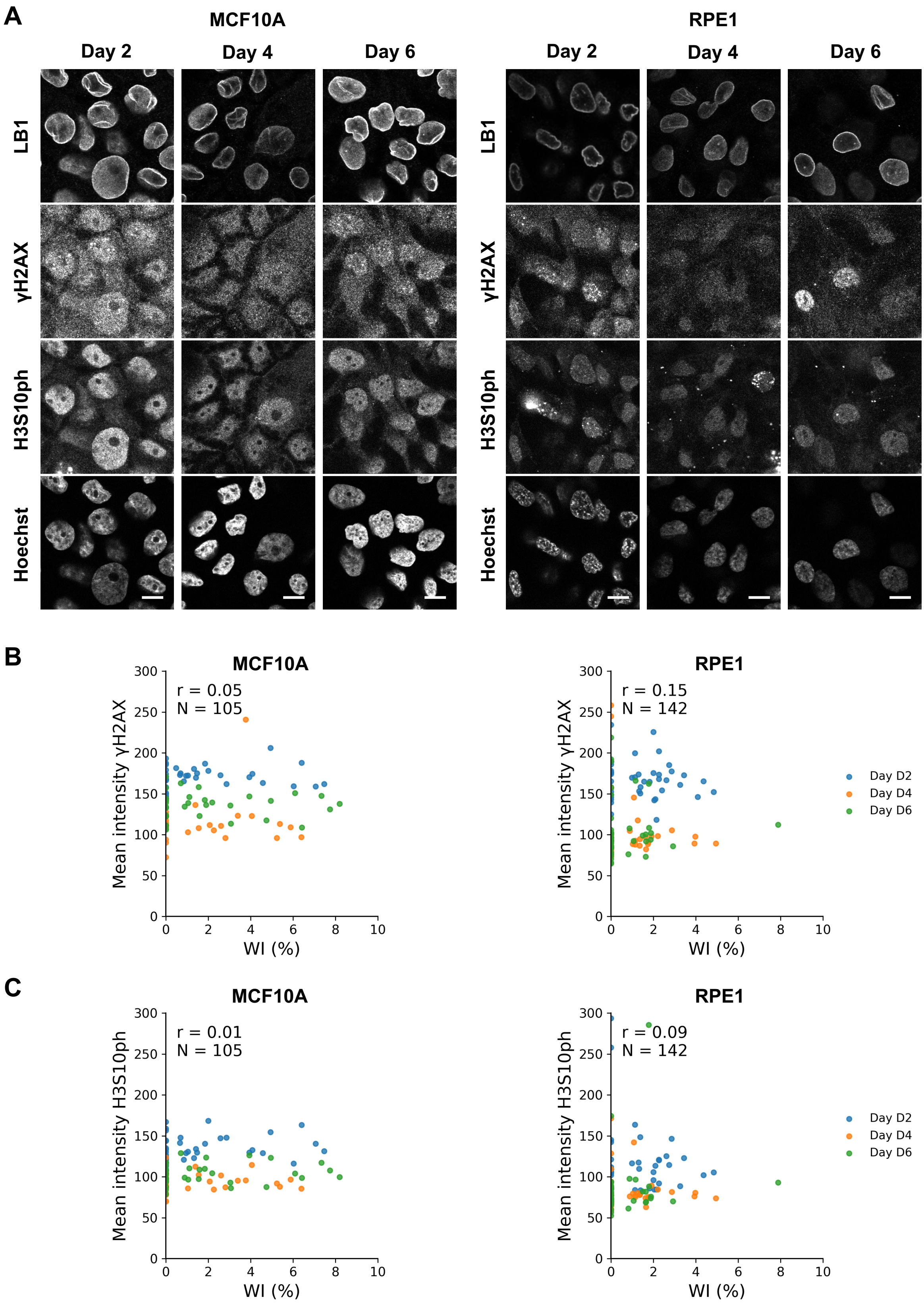


**Figure S11. Nuclear wrinkling is not associated with γH2AX or H3S10ph.**
**(A)** Representative immunofluorescence images of MCF10A and RPE1 cells stained for LB1, γH2AX, H3S10ph, and DNA (Hoechst). Single confocal sections are shown.
**(B, C)** Scatter plots showing the relationships between WI and mean nuclear intensities of γH2AX **(B)** and H3S10ph **(C)** in MCF10A (left) and RPE1 (right). Points represent single nuclei and are color-coded by culture day. Cells showing mitosis-associated H3S10ph were excluded. Pearson correlation coefficients (r) were calculated using pooled data from all days. n denotes the number of nuclei used for the analysis. There was little or no correlation between WI and γH2AX or H3S10ph intensity.
Scale bars, 10 µm.

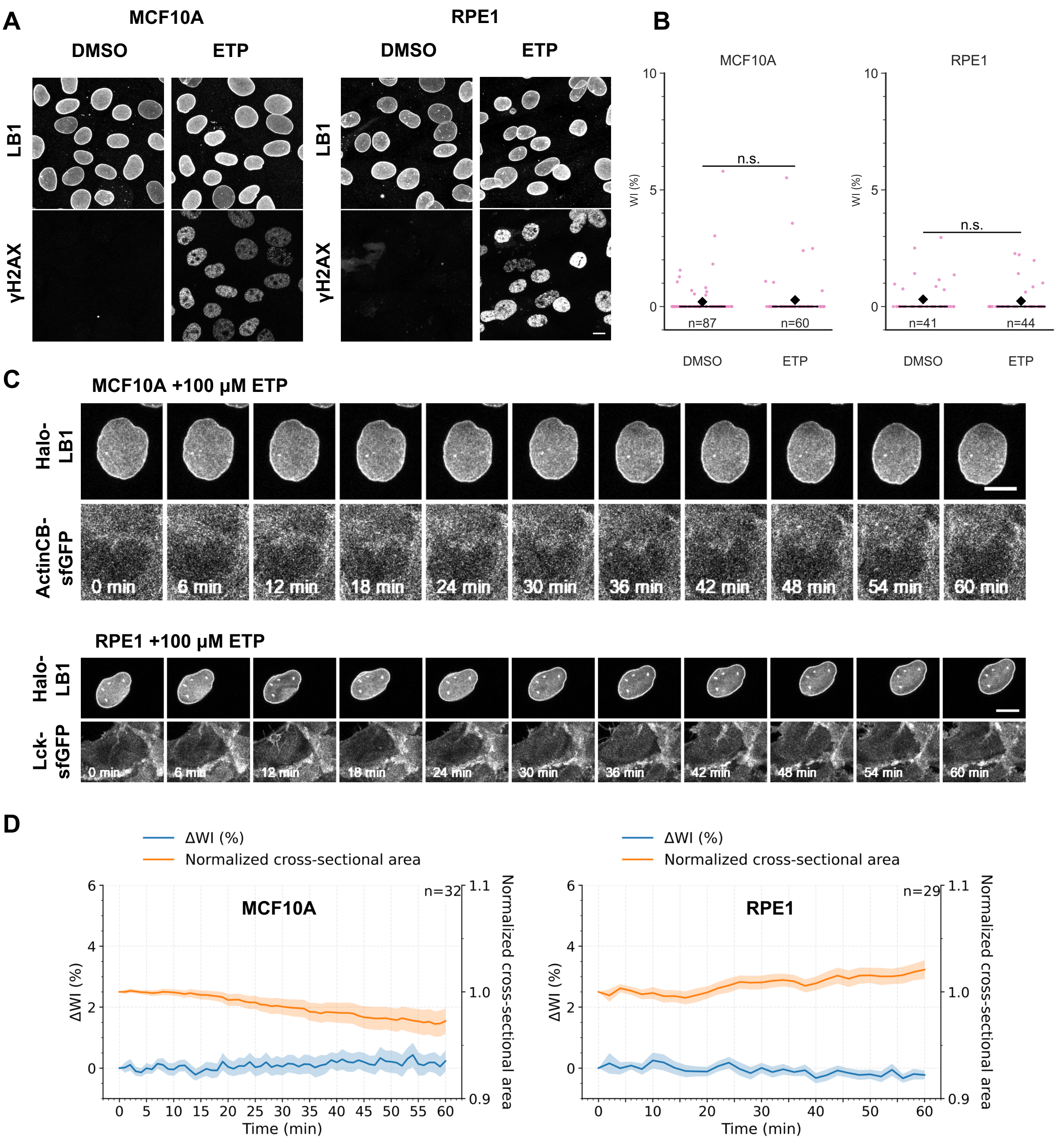


**Figure S12. DNA damage does not induce acute nuclear wrinkling.**
**(A)** Representative MIP images of MCF10A cells (left) and RPE1 cells (right) treated for 60 min with DMSO (control) or 100 µM etoposide (ETP) and stained for LB1 (top) and γH2AX (bottom).
**(B)** Quantification of the WI following ETP treatment. Data are presented as box plots with individual data points. Means (triangles) and total cell numbers (n), pooled from two biological replicates, are indicated. Statistical significance between DMSO and ETP treatment was assessed within each cell line (MCF10A and RPE1) using a two-sided Mann–Whitney U test. n.s., not significant.
**(C)** Representative time-lapse images of an MCF10A cell stably expressing Halo-LB1 and ActinCB-sfGFP (top) and an RPE1 cell stably expressing Halo-LB1 and Lck-sfGFP (bottom) following administration of 100 µM ETP at t = 0 min.
**(D)** Changes in WI relative to the first time point (ΔWI) and normalized nuclear cross-sectional area following ETP treatment in MCF10A (left) and RPE1 (right) cells. Data are presented as mean ± SEM (MCF10A, n = 32; RPE1, n = 29).
Scale bars, 10 µm.

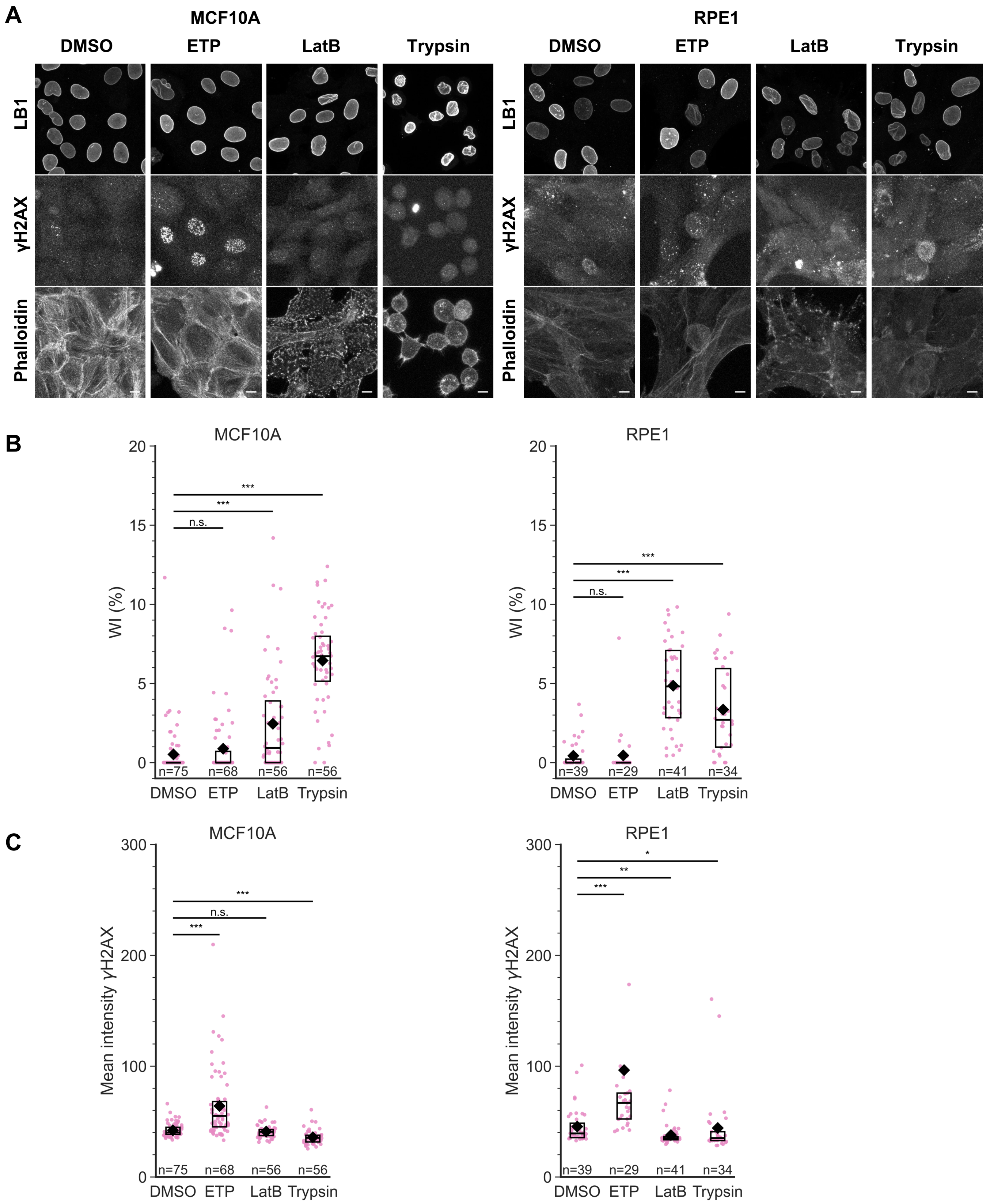


**Figure S13. LatB treatment and trypsinization do not induce detectable DNA damage.**
**(A)** Representative maximum-intensity projection immunofluorescence images of MCF10A and RPE1 cells treated for 15 min with DMSO (negative control), 100 μM ETP (positive control), 2 μM LatB, or 1 g/L trypsin, followed by staining for LB1, γH2AX, and F-actin (phalloidin). **(B, C)** Quantification of WI **(B)** and mean nuclear γH2AX intensity **(C)** following the indicated treatments. Data are presented as box plots with individual data points. MCF10A and RPE1 cells are shown in the left and right panels, respectively. Statistical differences among treatment groups were assessed separately within each cell line using the Kruskal–Wallis test followed by Dunn's multiple-comparisons test with Holm correction. Only comparisons between the DMSO control and each treatment are shown. *$p < 0.05$; **$p < 0.01$; ***$p < 0.001$; n.s., not significant. Data were pooled from two biological replicates and the total cell numbers (n) are indicated.

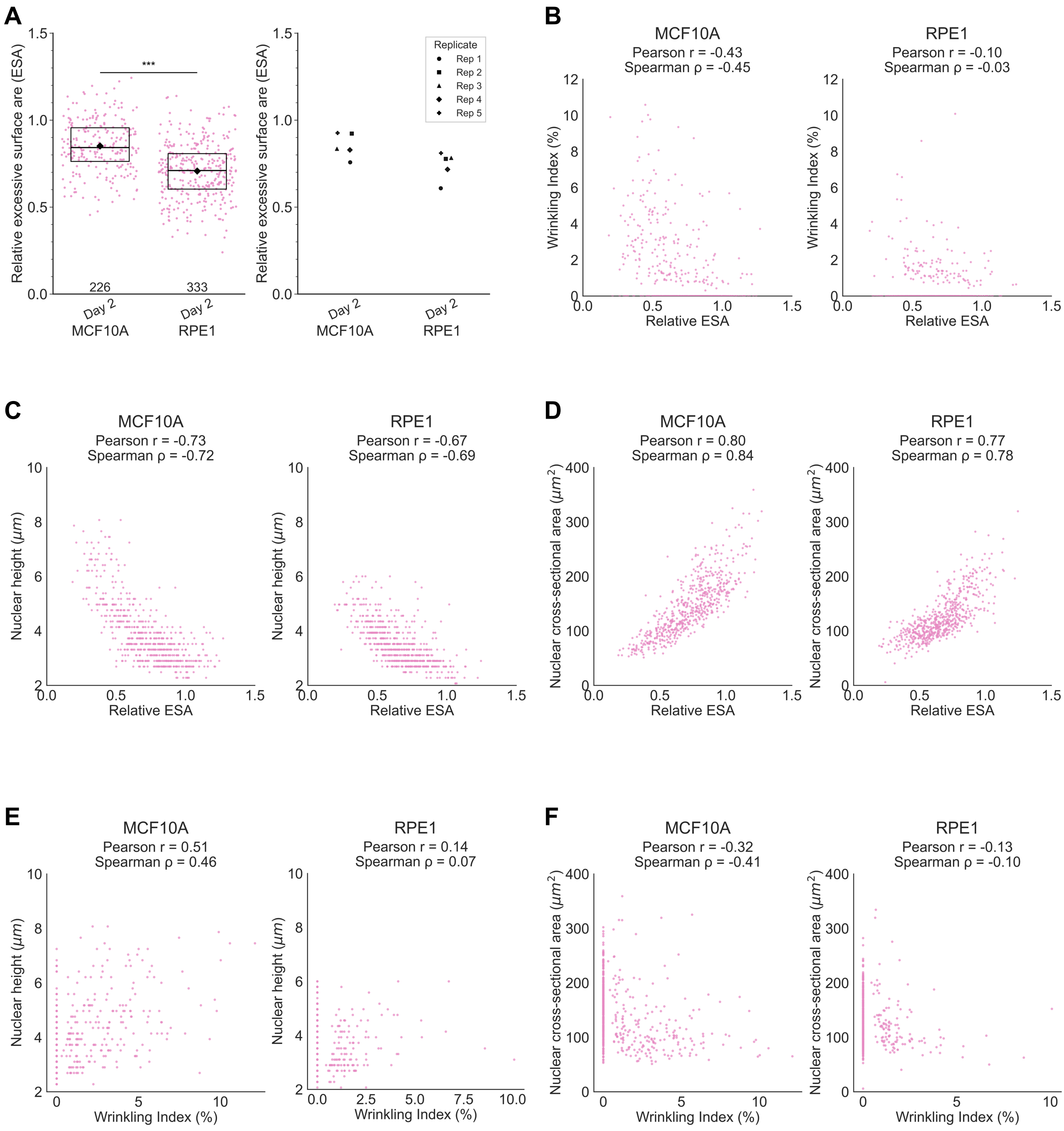


**Figure S14. Relative excess surface area is associated with nuclear wrinkling and nuclear morphology.**
**(A)** Quantification of the relative excess surface area (ESA) of taut nuclei (WI ≤ 0.1) at Day 2 in MCF10A and RPE1 cells. Left, ESA values of individual nuclei are shown as box plots with individual data points. Box plots indicate the median (line), interquartile range (box), whiskers extending to 1.5 × the interquartile range, and mean (diamond). Numbers of nuclei are shown below each group. Right, the mean ESA of each biological replicate is shown. Statistical significance between MCF10A and RPE1 was assessed using a two-sided Mann–Whitney U test. ***p < 0.001.
**(B–D)** Scatter plots showing the relationships between relative ESA and WI **(B)**, nuclear height **(C)**, and nuclear cross-sectional area **(D)** in MCF10A (left) and RPE1 (right). Data from Days 2–6 were pooled for each cell line (MCF10A, n = 575; RPE1, n = 586). Pearson's correlation coefficient (r) and Spearman's rank correlation coefficient (ρ) are indicated.
**(E, F)** Scatter plots showing the relationships between WI and nuclear height **(E)** and nuclear cross-sectional area **(F)** in MCF10A (left) and RPE1 (right) using the same dataset as in **(B–D)**.

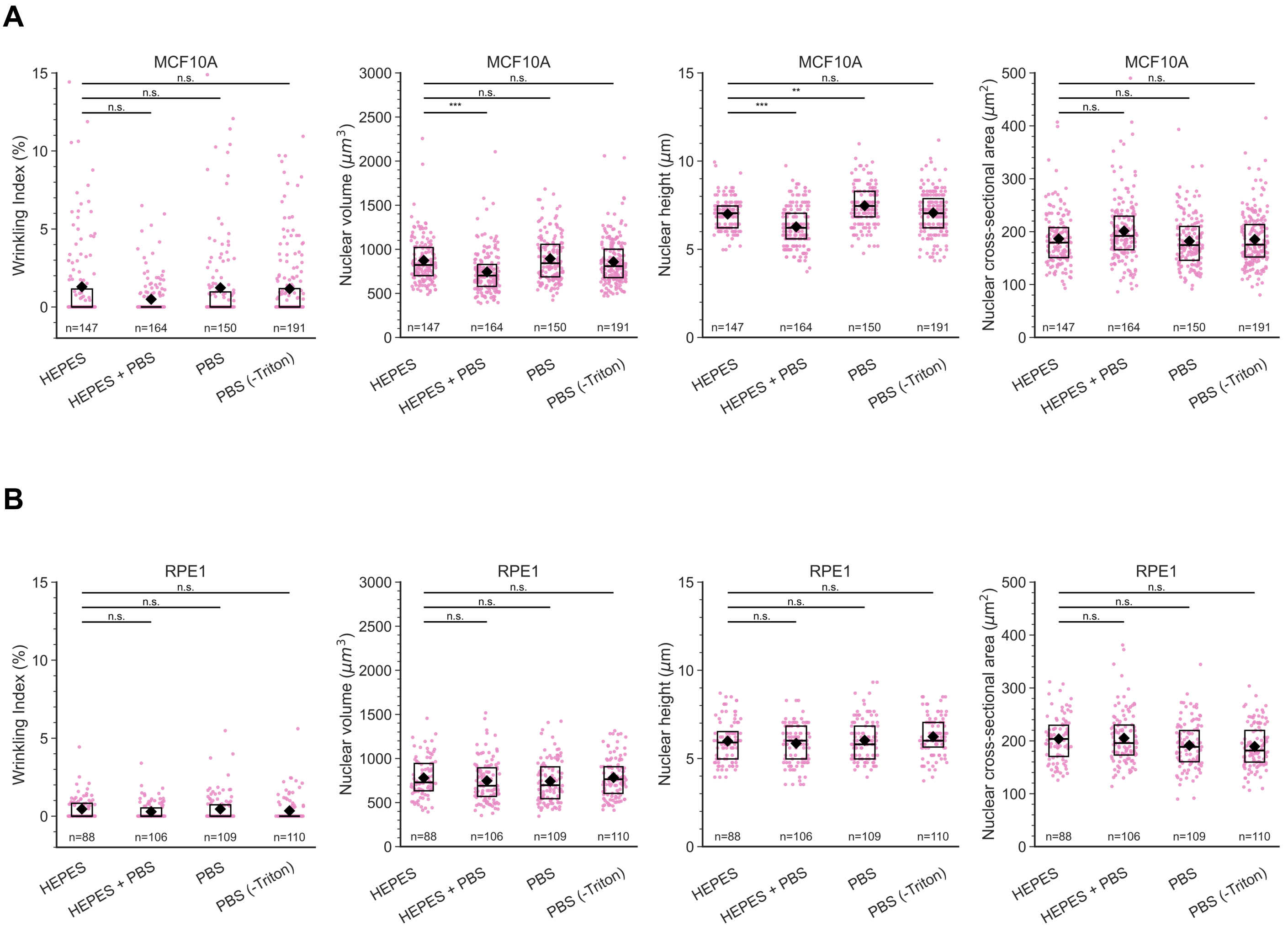


**Figure S15. Fixation conditions minimally affect nuclear geometry without altering the Wrinkling Index.**
MCF10A **(A)** and RPE1 **(B)** cells were fixed under four fixation conditions: **(1) HEPES**: 4% PFA, 250 mM HEPES-NaOH, and 0.1% Triton X-100 diluted in $H_2O$; **(2) HEPES + PBS:** 4% PFA, 250 mM HEPES-NaOH, and 0.1% Triton X-100 diluted in 1× PBS; **(3) PBS**: 4% PFA and 0.1% Triton X-100 diluted in 1× PBS; **(4) PBS (-Triton)**: 4% PFA diluted in 1× PBS. After staining for LB1 and DNA, Wrinkling Index (WI), nuclear volume, nuclear height, and nuclear cross-sectional area were quantified. The numbers of analyzed cells (n) are indicated. Data are presented as box plots with individual data points. In MCF10A, compared with the **HEPES** condition, the **HEPES + PBS** condition resulted in modest reductions in nuclear volume and nuclear height, whereas nuclear cross-sectional area and WI remained largely unchanged. In contrast, fixation conditions had little to no effect on nuclear geometry or WI in RPE1 cells. Statistical significance between the **HEPES** condition and each of the other fixation conditions was assessed within each cell line using the Kruskal–Wallis test followed by Dunn's multiple-comparisons test with Holm correction. $^{**}p < 0.01$; $^{***}p < 0.001$; n.s., not significant.
